# Spontaneous saccades reveal that human premotor cortex is involved in conceptual processing

**DOI:** 10.1101/2025.03.11.642662

**Authors:** Artur Pilacinski, Lia Wagner, Gabriel Besson, Maria Joao Matos, Eduardo Araujo, Christian Klaes

**Author notes:** **Corresponding author:** AP. Equal contribution.

## Abstract

Non-visual saccades (NVS) are spontaneous eye movements humans make while thinking. Their role is puzzling and there is no framework to explain them in terms of underlying neural systems. Here we studied neural activity preceding NVS when subjects performed standardized abstract thinking tasks. We used high-density EEG and focused on frontal and parietal channels representing oculomotor cortical areas. We found that NVS are distinctively preceded by neural activity changes in frontal and parietal EEG channels qualitatively different to that for voluntary saccades. However, only the ERP amplitude changes in frontal channels, but not in parietal channels, were different from saccadic preparation. A source reconstruction analysis additionally revealed that presaccadic neural activity originated mainly in the premotor cortex. Frequency spectrum analyses showed increased synchronization in alpha to low gamma bands suggesting complex top-down and bottom-up processing. We propose that this presaccadic activity while thinking represents evidence accumulation and attentional shifts, as shown before in the premotor cortex during sensorimotor tasks. Our findings suggest that human thinking appears to function similarly to human actions and engages premotor areas responsible for evaluating and manipulating objects also when evaluating and manipulating concepts.

## Introduction

What do apples and oranges have in common? What is similar between crime and punishment? Why do we pay taxes? Take a moment to answer these questions and you might notice that, before answering, you probably made a few eye movements as if in search of the right answer. It is a known but often ignored fact that such spontaneous, non-visual saccadic eye movements (NVS) accompany human thinking, that is mental task solving and mnemonic retrieval processes (Ehrlichman & Micic, 2012). Why they occur and what function they serve remains unknown. Here, we show that NVS are immediately preceded by specific activity changes in frontal regions including premotor cortex, indicating the crucial role of that area in conceptual operations (thinking).

Non-visual saccades remain a very understudied phenomenon and the existing research does not allow for building a coherent picture of their nature. Contrary to some opinions (Doherty-Sneddon and Phelps, 2005) NVS do not subserve others’ gaze aversion as they are also performed in darkness or without other people present (Ehrlichman & Barrett, 1983). Restriction of NVS does not seem to impact memory retrieval performance (Micic et al., 2010) suggesting they may be an epiphenomenon rather than a functional process (for an alternative view see: Scholz et al., 2016). They are, however, not fully suppressible and people move their eyes while thinking even when instructed to fixate gaze (Micic et al., 2010) indicating that NVS are automatically accompanying memory operations. On the other hand, non-visual saccades have been demonstrated to reflect mnemonic content organization such as spatial arrangement of items (Johansson et al., 2022; Damiano and Walther, 2019; Scholz et al., 2016) or the point in time (Hartmann et al., 2014; Martarelli et al., 2016). Although one could suspect these NVS to merely reflect visuospatial organization of memorized items, it comes as a surprise that substantially more saccades are produced when operating on verbal than on visual material (Weiner and Ehrlichmann, 1976). This indicates that NVS do not directly reflect reactivating the visual scene in memory but rather other processes such as activation of conceptual knowledge necessary for processing semantics.

Organization of conceptual knowledge has been long described as the core function of medial temporal lobe (MTL) (Bellmund et al., 2018; Constantinescu et al., 2016). Meanwhile, the main cortical centers controlling saccades span across posterior parietal (PPC) and premotor cortex (PM), comprising frontal eye fields (FEF), the main area of the brain responsible for programming voluntary eye movements (Munoz, 2002). While some known interactions exist between the MTL and eye movements systems, the mechanism that could possibly explain NVS remains unclear. For example, activity in MTL reflects integrating spatial information when exploring visual scenes (Leszczynski et al., 2024), about objects (Yeung et al., 2017), hippocampal maps are modulated by gaze direction (Meister et al., 2018) and parietal activity reflects that of the hippocampus (Whitlock et al., 2008). It is unclear, however, how retrieving mnemonic content could possibly initiate saccades. One recently suggested possibility is that hippocampus allows binding elements of the visual scene through receiving the corollary discharge accompanying visual exploration, thereby showing tight links between oculomotor system and memory encoding (Katz et al., 2022; Leszczynski et al., 2024). It is conceivable that such corollary discharge could then be reinstated too during memory retrieval, for example to enable shifting the focus of attention between elements of the conceptual space. While hypothetical, this perspective highlights the existing physiological links between medial temporal lobe and oculomotor regions in the primate brain that may be important for understanding NVS.

The core cortical regions for programming voluntary saccades are posterior parietal and prefrontal cortical areas (Pierrot-Deseilligny et al., 1995; Andersen & Buneo, 2002). Pre-saccadic planning activity was described earlier as predominantly visible in parietal EEG channels (Everling et al., 1997; Richards, 2003; Becker et al., 1972; Balaban & Weinstein, 1985). Posterior parietal cortex (PPC) has been proposed to integrate between memory and actions (Whitlock et al., 2008; Whitlock, 2017), and parietal oculomotor areas (Schluppeck et al., 2005) overlap with those involved in long term memory search (Berryhill et al., 2007; Vilberg and Rugg, 2008). PPC is also directly connected to superior colliculus (Pare and Wurtz, 1997), which suggests parietal activity may be triggering NVS as a byproduct of memory search processes. Whether it is the memory retrieval activity in parietal oculomotor areas that underlies NVS is unknown.

Posterior parietal cortex typically projects oculomotor programming signals to the frontal eye field, the main saccade-controlling area in the primate brain (Bruce et al., 1985). In humans, FEF is contained within the dorsal premotor cortex (Darby et al., 1996), indicating shared function of the two areas. Premotor cortex (PM) has been repeatedly demonstrated to be involved in a number of non-motor tasks, including working memory (Curtis et al., 2004; Curtis, 2006; Lindner 2010; Pilacinski et al., 2020), evaluating evidence during decision-making (Thura et al., 2012) and internal shifts of attention (Peelen et al., 2004). The premotor theory of attention further suggests that these internal shifts of attention reutilize saccade programming circuitry (Rizzolatti et al., 1996). Premotor cortex has also been shown to be involved in memory search (Kapur et al., 1995).

The organization of conceptual knowledge has long been suggested to reutilize sensorimotor computations, e.g. through representing cognitive spaces in a similar way as physical space (Tolman, 1948; Bellmund et al., 2018). In our view, non-visual saccades are a window into the evolution of human thinking by providing a link between the memory and the action systems in the human brain. The available evidence suggests that the frontal and parietal cortex may be involved in generation of NVS during memory search, although the particular involvement of each of these areas has not been explored. It is likewise unknown how neural mechanisms underlying NVS compare to typical programming of goal-directed saccades. Lastly, it is unclear whether NVS are a function or merely a byproduct of human thinking.

Here, we tackled these questions using high-density EEG, by investigating the neural activity immediately preceding non-visual saccades, while volunteers were solving abstract reasoning and knowledge tasks, with restricted and unrestricted gaze and with open and structured memory search conditions. In addition, we compared the presaccadic activity for NVS with voluntary saccades thereby controlling for unspecific preparatory signals such as readiness potential. Our central question was whether NVS are preceded by specific neural signals. If that was the case, we further reasoned that if NVS are a product of memory search mediated by posterior parietal cortex (as suggested by prior studies), then we should observe a neural activity change immediately preceding NVS in parietal channels. Conversely, if NVS were a product of attentional processing of conceptual information, they should be preceded by neural activity changes in frontal channels. Finally, if non-visual saccades arise from neural processing distinct to saccadic preparation, the neural activity changes preceding them should not be present before voluntary saccades.

## Methods

### Subjects

A total of 57 individuals (mean age 27.2 ± 8.1 years, 32 male, 5 left-handed) took part in the study. All participants had normal or corrected-to-normal vision with no history of neurological or psychiatric disorders. Participants were informed about the study procedures, were naive to the experimental hypothesis and received no financial compensation for their participation. Before participation, all subjects gave written informed consent in accordance with the Declaration of Helsinki and local ethical guidelines. The procedures were accepted by the local Ethics and Deontology Committee for Research of the Faculty of Psychology and Educational Sciences of the University of Coimbra. All datasets were anonymized before data collection began. Twelve participants were excluded from all analyses due to missing data or poor data quality as detailed in the Supplementary Materials (see Table S5), leaving an overall sample size of 45 participants (mean age 27.2 ± 8.0, 23 male, 4 left-handed) for the behavioral analysis. For the EEG analysis, further exclusion criteria based on EEG signal quality were applied, as described in the supplement, resulting in a final sample size of 29 participants (mean age 26.9 ± 7.0 years, 11 male, none left-handed). Due to a technical problem with data logging, there were just 12 subjects considered in analyses of the memory saccade control task (see Supplementary Materials, Table S5).

### Task

In the main experimental task, participants were supposed to answer forty questions taken from the Portuguese adaptation of Wechsler Adult Intelligence Scale-III (Wechsler, 1997). This task allowed us to have normalized stimuli, arranged progressively according to their difficulty and abstractness levels. The questions were presented verbally, through headphones (see Figure 1 A). The questions were split into “structured” and “open” according to the type of memory search they involved: 20 questions from the Similarities subtest were considered “structured”; 10 questions from the Information subtest, and 10 from the Comprehension subtest were considered “open”. The Information subtest evaluates long-term memory and general knowledge, e.g., “At what temperature does water boil under normal circumstances?” The Comprehension subtest assesses practical judgments, e.g., “Why do we pay taxes?” These were considered “open” questions due to the unrestricted memory search required. The Similarities subtest assesses abstract verbal reasoning by finding similarities between paired concepts, e.g., “How are an orange and a banana comparable?”. These were considered “structured” questions. Further examples can be found in the supplementary materials (see Table S1). Collectively, these subtests assess verbal conceptualization, vocabulary, abstract reasoning, categorical thinking, logical abstraction, and social intelligence, reflecting a broad range of cognitive functions and culturally acquired knowledge. The forty questions were divided into four blocks of 10 trials each, with randomized order within each block across subjects, and increasing abstraction across blocks. In each trial, a question was played through headphones and subjects were instructed to answer as best as they could. Correctness of answers was manually assessed according to the test instructions and saved in a binary form (correct/incorrect). The only displayed item during this task was a centrally-positioned bright fixation cross on a dark grey background. Subjects were randomly assigned to two groups: “fixed gaze” in which they were told to keep looking at the fixation cross and “free gaze” in which they were not instructed to do so.

**Figure 1.**
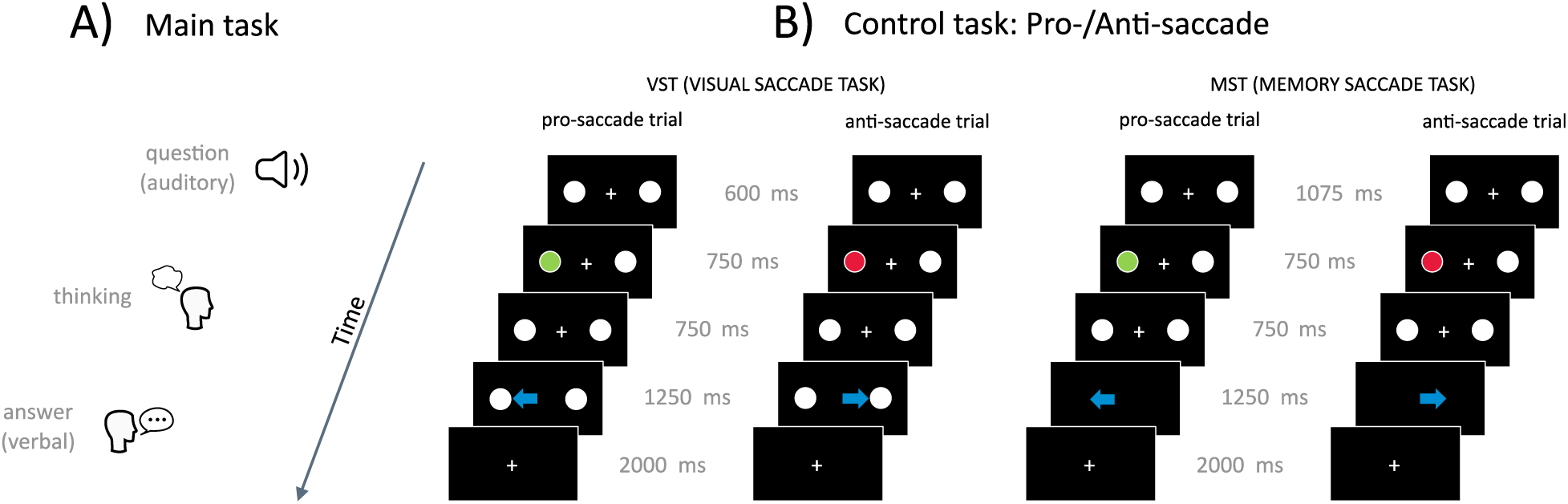
Experimental tasks timeline. A) Main task. The question was played through headphones; next, the participant was given time to think of an answer; after they responded, the next trial started. B) Control tasks: subjects were instructed to saccade to (green cue) or away from (red cue) the cued location (white dot). The saccade should be executed either towards the visible (VST) or memorized (MST) target. Blue arrows indicate saccade epoch. Objects not to scale.

A control task was used to assess goal-directed saccadic programming, as well as to differentiate between visual and memory saccade programming components (see Figure 1 B). The task was split into the following conditions: visual vs. memory saccades and pro- vs. antisaccades. In the visual condition, participants were instructed to make saccades either toward (prosaccade) or away from (antisaccade) a blinking dot located in the periphery. In the memory condition, participants were asked to make saccades to the location where a target dot had been before it disappeared. In both conditions, the saccade had to be executed after a delay (when the fixation cross disappeared).

### Eye Tracking

Eye movements were recorded at 100 Hz using the HTC VIVE Pro Eye, which has an embedded eye tracker in a VR head-mounted display and allows for accurate recording of eye data in immersive environments (Sipatchin et al., 2021). The VR environments were developed to serve as experimental tasks (see Figure 1 A, B), and an eye tracking algorithm was implemented in Unity with the use of SRanipal software development kit from HTC. The use of such a head-mounted display allows for reliable EEG recordings (Tauscher et al., 2019, including EEG systems similar to the one used here (Dejbara et al., 2019).

For the behavioral oculomotor analysis, saccade and blink frequencies, pupil diameter, saccade amplitudes, peak velocities, and angles were extracted for each trial and participant, focusing on the period after each question until the end of each trial (see Supplementary Methods). Parameters were averaged across combinations of factors for each participant, specifically blocks 1-4, condition type (fixed/free), and question type (open/structured). To assess eye parameters during the cognitive engagement, GLMM analyses were used as an alternative to the repeated measures ANOVA, with the fixed factors block (1-4), condition type (fixed/free) and question type (open/structured). Subjects were considered as the random effects grouping factor. GLMM analyses were performed separately for the dependent variable answer duration, saccade frequency, saccade amplitude, blink frequency and pupil average. Only ‘correct’ trials were included in the statistical analysis to minimize confounds in an already complex statistical model and ensure a clearer interpretation of cognitive engagement. The final trial count in the EEG analyses for the main tasks was 2884 (mean 100 ± 52.26 per participant), and for the control tasks was 2168 (mean 94 ± 11.95 per participant).

### EEG Recordings

EEG data was recorded using either a 64-electrode dry or a 256-electrode wet EEG system (eego mylab, ANT Neuro, Enschede, Netherlands), and event triggers were sent via the Lab Streaming Layer (LSL) protocol between the computers of the VR system and the EEG software (eego software, ANT Neuro), ensuring precise synchronization of the stimulus and eye tracking events within the EEG data. Initially, the 256-channel system was used for the first eleven participants, but due to technical issues, the 64-electrode system was used for the remaining eighteen participants. There were no statistically significant differences between both systems in terms of signal quality that would influence our results (see Supplementary Methods). Electrodes were arranged in an equidistant montage with the left and right mastoids as online ground and reference for the 64-channel setup, and CPz and the left mastoid for the 256-channel setup. Impedances were kept below 25 kΩ, and signals were recorded at 500 Hz. Preprocessing was done with EEGLAB and MNE-Python, including checks for sampling rate stability, application of a 1 Hz high-pass filter and 40 Hz low-pass filter, and removal of 25 Hz environmental noise using a notch filter. A 50 Hz notch filter removed remaining line noise. Flatline channels were removed using EEGLAB’s pop_clean_rawdata function, and noisy channels were rejected using pop_rejchan based on kurtosis, probability, and spectra, followed by manual inspection. Interpolated channels were re-referenced to the average of all electrodes. Eye movements were controlled using eye tracking data. EEG events were processed separately for the main and control tasks, and epochs were extracted from −200 to 100 ms relative to saccade onset. Saccades preceded by other saccades or blinks within 200 ms were excluded. A baseline correction from −200 to −170 ms was performed using single-trial normalization. We decided for this time window based on typical inter-saccadic latencies (Rayner et al., 1983). This allowed us to scrutinize signal changes immediately preceding NVS. Epochs were rejected if spikes were detected, and semiautomatic artifact rejection occurred for amplitudes exceeding 50 µV between samples or 100 µV over 50 ms. As the epochs were thus free of eye movements, ICA was not used to clean further the signal to avoid the risk of eliminating relevant data (e.g. saccade planning), particularly given recent evidence that automated ICA rejection, such as of eye movements (clearly present in their data), does not reliably increase the actual contrasts studied (Delorme, 2023). Four regions of interest (ROIs), frontal left (FL), frontal right (FR), parietal left (PL), and parietal right (PR), were defined, distributed around corresponding electrodes F3, F4, P3 and P4 in the 10-20 electrode system (see Figure 3 A and Figure S1 A, B). Since the FEF and PPC are the primary areas of interest in saccade research and are both superficial and spatially distinct, this setup allowed us to effectively distinguish neural activity in these regions.

### ERP analysis

For the EEG analysis, ERP analysis was performed using MNE-Python. An interval of −150 to −20 ms prior to saccade onset was used for statistical evaluation of presaccadic activity (Revankar et al., 2020). The mean amplitude of all epochs within this interval was calculated for each condition and participant. Potentials were then averaged across electrodes within each ROI, allowing for the assessment of neural dynamics associated with saccade preparation across different ROIs. Grand average ERPs were calculated for ERP visualization. For topographic visualizations of grand average ERPs, all electrodes from both EEG systems were included.

Statistical analysis of ERP data was performed either on each ROI or on combined frontal and parietal ROIs. A Kolmogorov-Smirnov test was used on averaged amplitudes to confirm normality prior to ANOVA analyses. Averaged EEG amplitudes of conditions were statistically compared using 4 × 2 or 2 × 2 repeated measures ANOVAs with ROI (4 ROIs and 2 ROIs, respectively) and condition type as factors. In cases of violation of sphericity (as indicated by Mauchly’s test), Huynh-Feldt corrected p-values were used. For post hoc analysis, paired t-tests were conducted with Holm correction for multiple testing. To investigate the temporal dynamics of signal divergence between main and control tasks, the same statistical approach was applied using a sliding window of 10 ms within the previously defined interval. Mean amplitudes were calculated within each window for each participant and condition, and paired t-tests with Holm correction were used to identify the time points at which significant differences emerged.

### Frequency spectrum analysis

Spectral decomposition was conducted using EEGLAB to assess neural activity in specific frequency bands associated with saccade preparation. The analysis focused on three frequency bands: alpha (8–12 Hz), beta (13–30 Hz), and low-gamma (30–45 Hz). Power spectral density (PSD) was computed for the interval of −150 to −20 ms prior to saccade onset. For each condition, PSD was averaged over time and the selected ROIs. Statistical analysis was performed separately for each frequency band on combined frontal and parietal ROIs. Repeated measures ANOVAs were conducted with ROI and task type as factors to evaluate differences in PSD values. In cases of violation of sphericity (as indicated by Mauchly’s test), Huynh-Feldt corrected p-values were used. For post hoc analysis, paired t-tests were conducted with Holm correction for multiple testing.

### Source localization

The forward model and the ERP for each condition were used to estimate an inverse model using the standardized low-resolution brain electromagnetic tomography (sLORETA) method (Pascual-Marqui, 2002). We extracted source estimates for each condition and computed spatial adjacency in the source space to enable cluster-based permutation testing. We performed a non-parametric, two-tailed cluster-based permutation test, involving voxel-wise comparisons to zero and correcting for multiple testing. We assessed statistical significance using a p-value threshold of 0.05. We projected t-values into sLORETA images to visualize source activity, highlighting regions with significant neural dynamics. We obtained anatomical labels using the FreeSurfer parcellation scheme to map source activity to specific cortical regions, focusing on cortical involvement in saccadic movements. Subcortical areas were not captured due to parcellation limitations. We aligned source estimates with these labels to investigate neural activity in distinct areas. For each significant cluster identified, we extracted the maximum t-value for each label.

## Results

### Behavioral and oculomotor results

First, a decrease in percentage of known answers can be observed across blocks for structured questions while open questions show a significant drop in block 2 with no overall decrease across blocks (see Figure 2 A). Additionally, interactions between question type and condition type revealed fewer known answers for the fixed gaze condition for structured questions only (see Figure 2 B). A significant main effect was found of the factor block (*χ^2^* = *53*.*151*, *p* < *0*.*001*), indicating that performance dropped as abstractness increased. No significant main effect was found for question type (*χ^2^*= *3*.*398*, *p* = *0*.*065*) or condition type (*χ^2^* = *2*.*702*, *p* = *1*.*000*), indicating that the percentage of known answers did not reliably differ between structured and open questions or between fixed and free gaze conditions. Significant interaction effects were found for block × question type (*χ^2^*= *112*.*722*, *p* < *0*.*001*) and question type × condition type (*χ^2^*= *7*.*987*, *p* = *0*.*005*), suggesting that the effect of block varied across different question types. Post hoc Wilcoxon signed-rank tests revealed a significant difference for block 1 and block 2, block 1 and 4, and block 3 and block 4 (*p* < *0*.*001*). Additionally, a significant difference was found between block 1 and block 3 (*p* = *0*.*031*). Post hoc Wilcoxon rank-sum tests also showed a difference between fixed and free gaze condition (*p* = *0*.*029*). For interactions between question type and condition type (Figure 2 B), post hoc Wilcoxon rank-sum tests confirmed a significant difference between open compared to structured questions for the fixed gaze condition only (*p* = *0*.*026*), but not for the free gaze condition (*p* = *1*.*000*).

**Figure 2.**
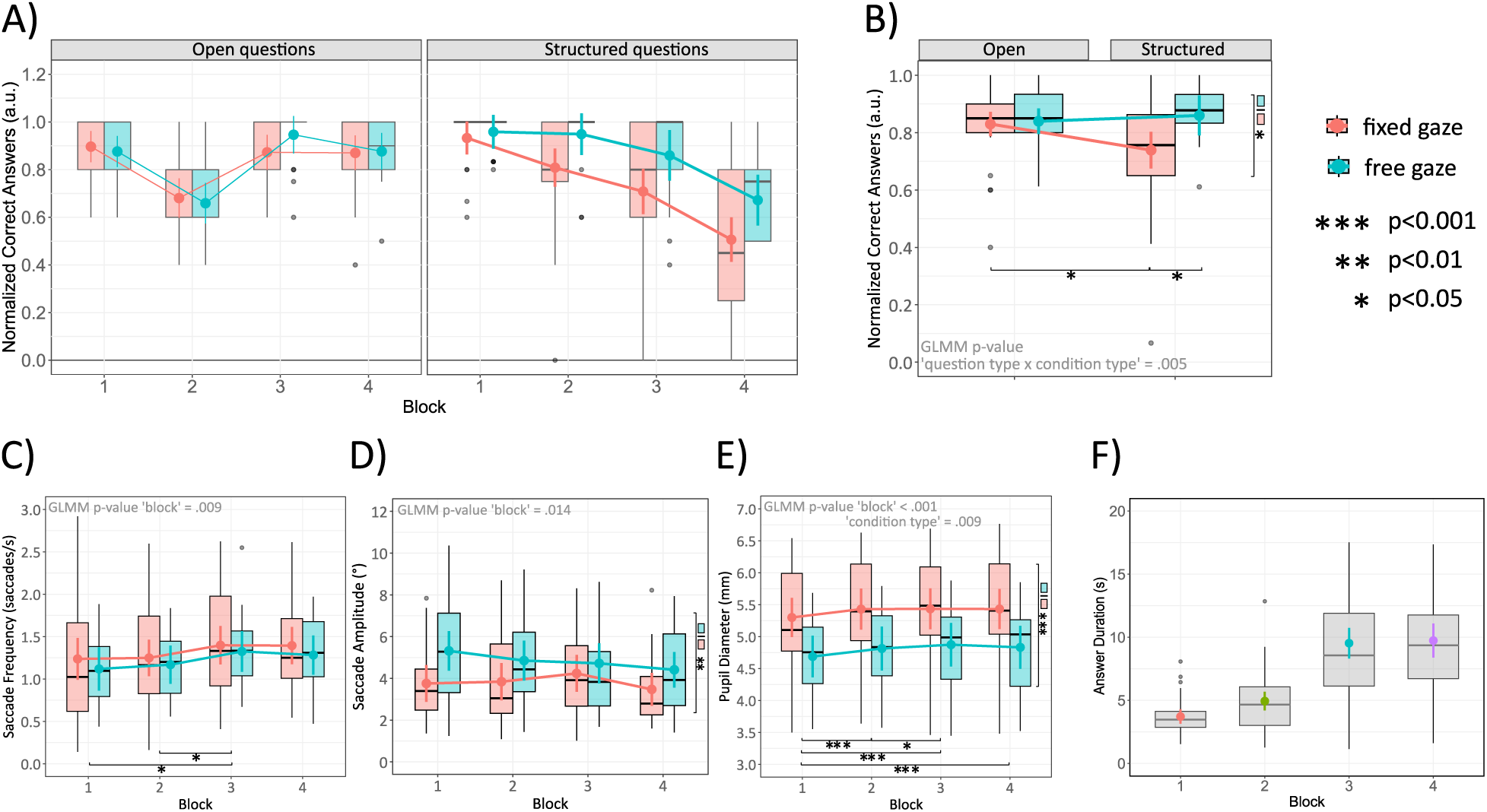
Behavioral metrics for different gaze conditions across fixed effects variables (n = 45). Box plots show the IQR with medians as horizontal lines. Mean values with 95% confidence intervals are marked as colored points, and outliers as individual points. Only known response trials were included. A) Normalized correct responses are shown across blocks and B) different question types. C) Saccade frequency, D) saccade amplitude, E) pupil diameter and F) answer durations are shown across blocks. Error bars denote upper and lower quartiles, boxes denote interquartile ranges, solid horizontal bars denote medians.

Saccade frequency for condition type and question type remains approximately the same, but increases over blocks for both fixed and free gaze conditions (see Figure 2 C). For saccade frequency, a significant main effect was found of the factor block (*χ^2^* = *11*.*500*, *p* = *0*.*009*). Post hoc Wilcoxon signed-rank tests revealed a significant difference between block 1 and 3 (*p* = *0*.*017*) and block 2 and 3 (*p* = *0*.*043*). However, no significant difference was observed for block 4, including comparisons between block 1 and block 4 (*p* = *0*.*055*), showing that while we see an increase in saccade frequency in block 3, the frequency is slightly reduced again in block 4. The post hoc Wilcoxon rank-sum test for gaze condition type showed no significant difference (*p* = *0*.*604*).

Saccadic amplitudes slightly decreased across blocks (see Figure 2 D). For different question types, saccadic amplitude remains approximately the same, whereas in the fixed condition, mean saccadic amplitudes are decreased compared to the free gaze condition. For saccadic amplitude, a significant main effect of the factor block was found (*χ^2^*= *10*.*563*, *p* = *0*.*014*). Also, a significant interaction effect of block × question type × condition type (*χ^2^* = *9*.*784*, *p* = *0*.*020*) suggests that the effect of blocks varied across different question types as well as condition types. Regarding condition type, the post hoc Wilcoxon rank-sum test revealed a significant difference between fixed and free gaze conditions (*p* = *0*.*006*).

An increased pupil diameter as an indicator of cognitive load can be identified across blocks and in the fixed gaze condition compared to the free gaze condition (see Figure 2 E). Additionally, a slight increase in pupil diameter can be noted for open questions compared to structured questions. For pupil diameter, a significant main effect of the factor block (*χ^2^*= *24*.*720*, *p* < *0*.*001*), question type (*χ^2^* = *16*.*897*, *p* < *0*.*001*), and condition type (*χ^2^* = *6*.*750*, *p* = *0*.*009*) was found. Additionally, a significant interaction between block × question type was identified (*χ^2^* = *21*.*756*, *p* < *0*.*001*). Post hoc Wilcoxon signed-rank tests revealed a significant difference for block 1 compared to all other blocks (*p* < *0*.*001*). A significant difference was also found between block 2 and 3 (*p* = *0*.*029*). No significant differences were found between block 2 and 4 (*p* = *0*.*194*) or between block 3 and 4 (*p* = *0*.*311*). These results suggest that pupil diameter increased across the initial blocks, indicating rising cognitive load or engagement levels, but slightly decreased again in block 4.

Lastly, for answer duration (see Figure 2 F), a significant main effect was found of the factor block (*χ^2^*= *58*.*375*, *p* < *0*.*001*) and question type (*χ^2^* = *6*.*074*, *p* = *0*.*014*). There was also a significant interaction effect of block × question type (*χ^2^* = *24*.*804*, *p* < *0*.*001*), suggesting that the effect of blocks varied across different question types. Post hoc Wilcoxon signed-rank tests revealed a significant difference between block 1 and 2 (*p* = *0*.*002*), and all other blocks (*p* < *0*.*001*), except block 3 and 4, where no significant difference was found (*p* = *0*.*697*). The post hoc Wilcoxon rank-sum test for condition type showed no significant difference (*p* = *0*.*153*). This indicates that participants took longer to respond for more abstract questions.

Presaccadic potentials were analyzed for main versus control tasks, with the control task including both visual and memory saccade tasks. The analyzed interval (−200 to 50 ms) is referenced within the overall signal (c.f. Figure S6 A), as determined based on typical saccadic latency time (Rayner et al., 1983) to avoid intervening saccades. Within-condition baseline reference means that the reported effects reflect relative changes rather than absolute increases or decreases in activity. In the specified interval, the main task is characterized by a presaccadic positive frontal and negative parietal potential, while the control task shows a positive parietal potential and a smaller frontal potential in contrast to the main task (see Figure 3 A). Potentials peak at approximately 20 ms prior to saccade onset. At saccade onset, a negative frontal and positive parietal spike can be observed for both task conditions. Averaged presaccadic amplitudes are larger for the main task in frontal ROIs and smaller for the main task in parietal ROIs compared to the control task. Supplementary analyses confirmed that the observed ERP differences between main and control tasks are not driven by differences in saccade velocity or amplitude (see Supplementary Results and Figure S5). A larger polarization is also observed for frontal and parietal ROIs for the main task compared to the control task, as confirmed in the topographical visualization. Topographic maps illustrate the evolving spatial distribution of ERPs leading up to saccade onset (see Figure 3 E). As already indicated by the ERPs, the main task shows a strong positive frontal and negative parietal potential developing over time, peaking just before the saccade. In contrast, the control task exhibits a prominent positive potential in the parietal regions, which extends into the posterior frontal regions. However, overall, the frontal electrodes show less pronounced, negative potential. A larger cluster of activity is visible over the parietal electrodes. This pattern is congruent with previously reported presaccadic EEG activity (Everling et al., 1997; Richards, 2003; Becker et al., 1972; Balaban & Weinstein, 1985). To quantify these activity changes, we used repeated measures ANOVA using ERP amplitudes averaged over the presaccadic time window (−150 to −20 ms), with “task” (main/control) and “ROI” (frontal/parietal) as factors. This analysis showed significantly higher activity change preceding saccades during the thinking task in both channel groups (see Figure 3 B). A significant main effect of “task” (*F*(*1*, *28*) = *16*.*205*, *p* < *0*.*001*, *ω^2^*= *0*.*137*) and “ROI” (*F*(*1*,*28*) = *5*.*528*, *p* = *0*.*026*, *ω^2^*= *0*.*114*) was found with a large and medium to large effect size, respectively. Additionally, a significant interaction effect of “task” × “ROI“ (*F*(*1*,*28*) = *25*.*496*, *p* < *0*.*001*, *ω^2^*= *0*.*411*) with a large effect size was observed. Post hoc paired t-tests revealed a significant difference between main and control tasks for the frontal channels (*p* < *0*.*001*), as well as for the parietal channels (*p* = *0*.*003*). Significant differences were also identified for the main task between frontal and parietal ROI (*p* < *0*.*001*), while no significant difference was found for the control task between frontal and parietal ROI (*p* = *0*.*130*).

**Figure 3.**
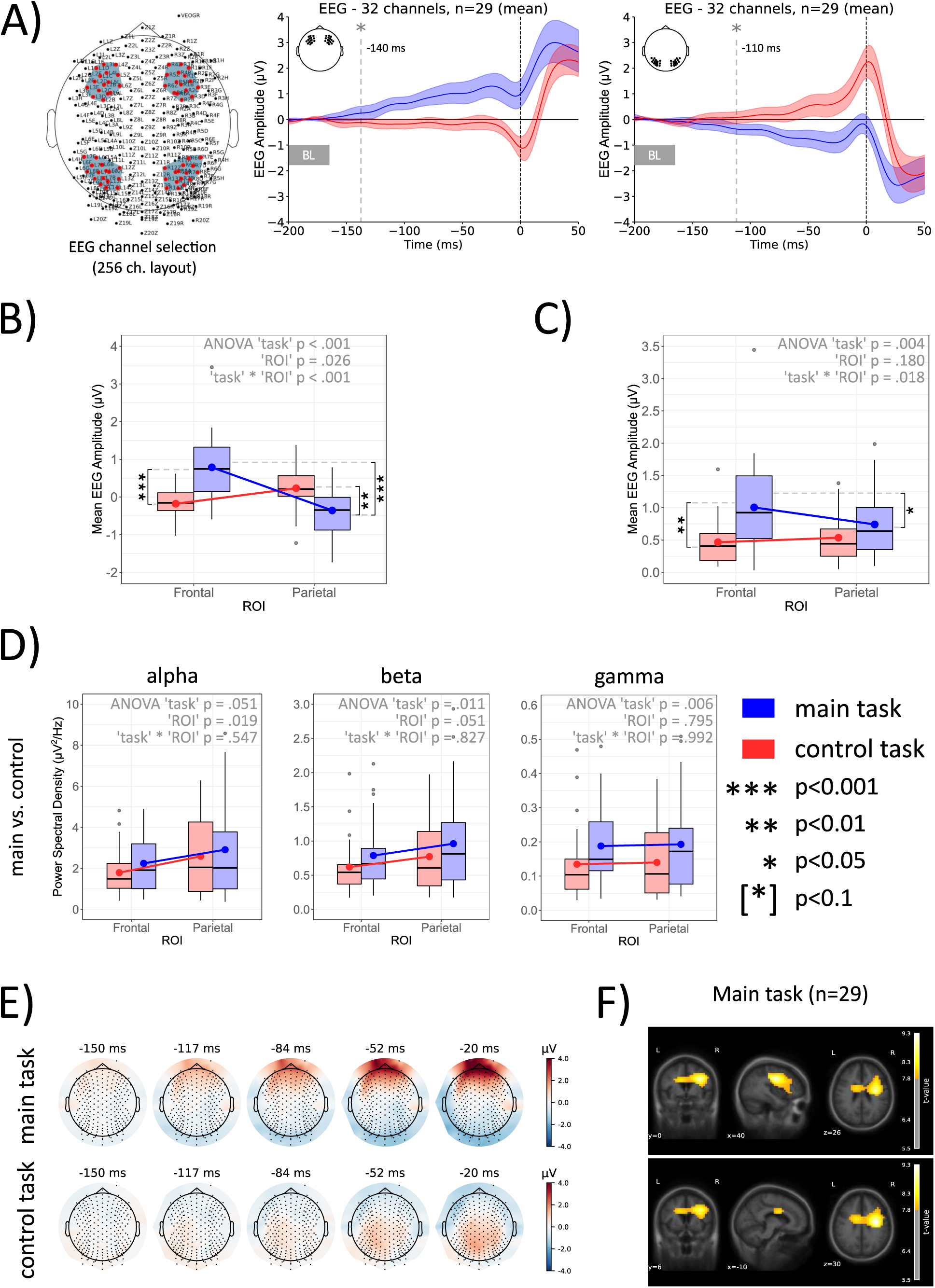
EEG results for main vs. visual and memory control tasks. A) ROI channels selection and ERPs relative to saccade onset for main and the control tasks; dashed lines indicate significant amplitude difference between ERP signals in main and control tasks; grey box denotes the baseline period; shaded areas denote 95% confidence intervals; B) Mean ERP amplitudes across tasks. C) Absolute values of ERP amplitudes across conditions. D) Modulation of alpha, beta and gamma frequency across tasks. E) Topographic maps of presaccadic activity across main and control tasks. F) Source localization. Error bars denote upper and lower quartiles, boxes denote interquartile ranges, solid horizontal bars denote medians.

As previous analyses showed that the main and the control tasks had distinct neural components, we next analyzed the absolute magnitude of pre-saccadic EEG activity changes (−150 to −20 ms) independent of potentially confounding effects of signal polarity (see Figure 3 C). We compared the absolute values of ERP amplitudes by conducting repeated measures ANOVA with “task” (main/control) and “ROI” (frontal/parietal) as factors. A significant main effect of “task” (*F*(*1*, *28*) = *10*.*087*, *p* = *0*.*004*, *ω^2^* = *0*.*145*) was found with a large effect size. No significant main effect was found for “ROI” (*F*(*1*,*28*) = *1*.*887*, *p* = *0*.*180*, *ω^2^*= *0*.*009*). Additionally, a significant interaction effect of “task” × “ROI“ (*F*(*1*,*28*) = *6*.*279*, *p* = *0*.*018*, *ω^2^*= *0*.*047*) with a small effect size was observed. Post hoc paired t-tests revealed a significant difference between main and control tasks for the frontal channels (*p* < *0*.*001*), but no significant difference for the parietal channels (*p* = *0*.*265*). Significant differences were also identified for the main task between frontal and parietal ROI (*p* = *0*.*035*), while no significant difference was found for the control task between frontal and parietal ROI (*p* = *0*.*477*). These analyses show that absolute signal amplitude changes in prefrontal channels in the main task were greater than those in parietal channels and when compared to the saccadic preparatory activity in the control tasks. The sliding window analysis revealed the time points at which ERP signals significantly separated between the main and control tasks in the frontal and parietal ROIs (see Figure 3 A). In the frontal ROI, signal separation was first observed at −140 ms prior to saccade onset (*p* = *0*.*038*). In the parietal ROI, signal separation occurred later, at −110 ms prior to saccade onset (*p* = *0*.*042*). The results demonstrate a 30 ms difference between the onset of significant signal change in the frontal vs. parietal ROIs, with the frontal onset preceding the parietal one. This temporal difference underscores the distinct timing of neural processes involved in NVS across these regions.

In the next step, we analyzed power spectral density for alpha, beta, and low-gamma bands (see Figure 3 D). For each band separately, we conducted an ANOVA with factors “Task” and “ROI”. These ANOVAs showed a significant increase in power spectral density in the main task due to a significant “Task” main effect for beta (*F*(*1*, *28*) = *7*.*414*, *p* = *0*.*011*, *ω^2^*= *0*.*029*) and gamma bands (*F*(*1*, *28*) = *9*.*016*, *p* = *0*.*006*, *ω^2^* = *0*.*058*). There was also a nearly-significant main effect of Task in alpha band (*F*(*1*, *28*) = *4*.*14*, *p* = *0*.*051*, *ω^2^* = *0*.*011*). There was a significant ROI main effect in alpha (*F*(*1*, *28*) = *6*.*171*, *p* = *0*.*019*, *ω^2^* = *0*.*036*) and a near significant ROI main effect in beta bands (*F*(*1*, *28*) = *4*.*174*, *p* = *0*.*051*, *ω^2^*= *0*.*020*). No Task x ROI interaction effect was significant (*p* > *0*.*5*).

Lastly, source localization revealed strong, focused neural activity primarily in the right hemisphere. Figure 3 F shows clusters in the frontal regions, with the peak activation in the right superior and middle frontal gyrus (*peak t* = *9*.*3*). Activity extends posteriorly into the motor cortex and precentral gyrus, and medially, involving orbitofrontal and medial areas. Notable activation clusters was also found bilaterally in the superior frontal cortex (*t* = *8*.*17*), precentral and postcentral regions (*t* = *8*.*21*), the insula (*t* = *8*.*14*), and rostral anterior cingulate (*t* = *7*.*98*). Additional activation clusters, though less intense, occurred in the superior parietal cortex (*t* = *7*.*95*), medial orbitofrontal cortex (*t* = *7*.*88*), and precuneus (*t* = *7*.*97*). This pattern was different in control tasks (see Supplementary Results).

## Discussion

Non-visual saccades are a not well understood phenomenon accompanying conceptual operations. Here, we show that NVS are directly preceded by neural activity changes in frontal EEG channels. The source localization analysis additionally indicated that the pre-NVS activity originates in the premotor cortex. These pre-NVS changes are qualitatively different from signals preceding voluntary saccades indicating that NVS are driven by other processes than oculomotor preparation. While pre-NVS activity changes are seen in both frontal and posterior channels, only the activity change in frontal channels significantly differ from that preceding voluntary saccades. As we used abstract conceptual tasks, we can determine that neural activity before NVS is not merely reflecting mnemonic processing or retrieval of visuospatial content.

It has long been a subject of dispute in philosophy, psychology, and neuroscience whether concepts are represented in pictorial/visuospatial or verbal/abstract form (see e.g., Pylyshyn, 2003). Edward Tolman proposed that past experience and spatial cognition are arranged in the form of spatially organized maps (Tolman, 1948). This theory implied that the knowledge map builds on representations of space, and the two likely share neural components. Since then, it has been confirmed that spatial maps are organized by the hippocampal-entorhinal system (O’Keefe & Dostrovsky, 1971) using a grid-like code (Hafting et al., 2005), and that this system also organizes conceptual knowledge using similar principles (Constantinescu et al., 2016). However, just as spatial maps need to be translated into neural codes understood by the motor system, stored conceptual knowledge needs to be processed for daily operations. One recently suggested possibility is that the hippocampus plays a crucial role in binding elements of a visual scene by receiving corollary discharge signals that accompany visual exploration. By receiving these signals, the hippocampus may be able to temporally align incoming visual information with the sequence of eye movements, thereby supporting the formation of coherent scene representations in memory (Katz et al., 2022; Leszczynski et al., 2024). It is conceivable that these same corollary discharge signals are reinstated during memory retrieval to recreate aspects of the original exploratory state, thereby enabling attention to be internally shifted between different components of the remembered scene or conceptual space. This possibility aligns with models of memory that propose dynamic and reconstructive processes, where internally generated signals guide the reactivation of stored representations. In this context, the MTL may contribute not only to encoding but also to the active reconstruction of memory traces via simulated sensory-motor interactions.

It has been proposed that the parietal lobe acts as an interface connecting the MTL memory system with areas responsible for actions (Whitlock, 2017). The main role of the PPC seems to lie in integrating representations of world-centered space into body-centered maps used for locomotion and actions (Whitlock et al., 2008). In the cognitive domain, there is also strong evidence showing the parietal lobe’s involvement in long-term memory search but not in the recognition of remembered objects, indicating that the parietal cortex is not merely involved in storage but rather in the active exploration of the knowledge space (Cabeza et al., 2008). Motor (and oculomotor) areas in the posterior parietal cortex share strong reciprocal connections with the premotor cortex (Pandya & Kuypers, 1969), as both regions perform complementary roles in controlling actions (Curtis, 2006; Cisek, 2007; Westendorff et al., 2010; Pilacinski and Lindner, 2019). This tight fronto-parietal interrelation in processing actions seems to be retained when it comes to operating on memory. Both PM and PPC seem to process different stages of search and retrieval of mnemonic content, with PM controlling memory search and PPC successful retrieval of information (Kapur et al., 1995). Attentional shifts underlying memory search were demonstrated to increase functional connectivity between PM and the parietal cortex, suggesting a tight information exchange in the process of retrieving and manipulating mnemonic information, which relies critically on selecting relevant and suppressing irrelevant information (Heinen et al., 2017). But why would the premotor cortex be involved in thinking? The answer lies in how it is involved in actions.

The primary function of the premotor cortex seems to be its ability to represent complex, coordinated multi-joint movements (Graziano and Aflalo, 2007) such as those needed for programming hand trajectories or gestures (Pilacinski and Lindner, 2019; Wong et al., 2019), or handwriting (Kadmon Harpaz et al., 2014). The premotor cortex appears to process both motor and semantic levels (i.e., higher-order goals) of actions (Rizzolatti et al., 1996) and is critically involved in executing relevant and suppressing irrelevant actions from among options represented by the parietal cortex (Cisek, 2007; Lindner et al., 2010; Pilacinski and Lindner, 2019). Neurophysiological data further suggest that the key function of the premotor cortex in programming movements is selecting from among competing action plans represented by the posterior parietal cortex, using weighting input from more frontal areas (Cisek, 2007). The tight connectivity between PM and PPC further suggests that this inter-areal exchange facilitates the promotion of relevant and suppression of irrelevant information, e.g., actions. In conceptual processing, such reciprocal exchange between PM and PPC may be utilized in a similar way, in which frontal areas (mediated by PM) select between competing information retrieved from long-term memory by parietal areas through top-down modulation, as suggested by ERP dynamics. This idea seems to be further supported by spectral analysis data, showing that immediately before each NVS, we observe a substantial increase in all three EEG bands we measured, indicating a complex computational process required by top-down and bottom-up modulation of cognitive content (Karch et al., 2016; Strube et al., 2021). These distinctive patterns of activity may reflect the evaluation of conceptual information by PM in a way similar to that during sensorimotor decision-making (Cisek et al., 2009; Thura and Cisek, 2014).

We believe that the gradual change of activity before NVS likely represents neural computations required for evaluating conceptual information, similar to evidence accumulation during sensorimotor decision-making (Cisek et al., 2009; Thura and Cisek, 2014). The subsequent saccade could then occur due to the shifting of attention to another sub-element of the currently-processed part of conceptual space. It has been shown that covert shifts of attention indeed result in involuntary eye movements, likely due to direct connections between frontal eye fields and the superior colliculus (Peterson et al., 2004). Complex conceptual operations require multiple such shifts of attention between concepts or their elements, and these shifts are likely to underlie NVS (Scholz et al., 2017) in a way similar to how saccades scan the visual space, identifying its (semantic) components. The premotor theory of attention proposes that covert attentional shifts and saccade planning are identical or closely-related processes involving mainly premotor cortex (Rizzolatti et al., 1987; Smith and Schenk, 2012). Our results suggest that non-visual saccades arise from specific processing within the premotor cortex, distinct from mechanisms involved in voluntary saccade planning. This interpretation is further supported by the fact that pre-NVS activity remained consistent even when subjects were instructed to withhold eye movements, indicating that non-visual saccades are preceded by a dedicated neural substrate. Nonetheless, the fact that neural activity preceding NVS seems to originate in the premotor cortex, a common neuroanatomical substrate between NVS and voluntary saccades, may explain why mental operations on concepts trigger NVS. As frontal eye fields directly project to the superior colliculus, it is likely that mental operations involving premotor computations while processing concepts may collaterally initiate eye movements as a byproduct of thinking. The fact that restricting gaze did not substantially worsen mnemonic performance supports the notion that NVS are an epiphenomenon rather than a functional component of conceptual processing.

## Limitations

Our study focuses on cortical processing responsible for thinking, considering the role of frontal and parietal areas in conceptual processing triggering NVS. Yet, human thinking involves numerous subcortical structures such as the hippocampus or basal ganglia, which are critically involved in information storage and selection. Due to EEG limitations, we cannot address the role these subcortical structures play in processing resulting in NVS. Although posterior parietal cortex and premotor cortex can directly exchange information through a one-synapse connection (Pandya & Kuypers, 1969), theoretically allowing for a direct exchange of information, we believe there may be substantial involvement of subcortical structures in how conceptual information is processed (Cisek et al., 2009). Due to EEG’s poor spatial resolution, our reasoning about the premotor cortex is mainly based on assumptions given its role in eye movements and sensorimotor processing rather than direct functional localization. We do acknowledge, however, that conceptual operations involve a long cascade of processes including emotional evaluation, evidence weighting, and everything that constitutes individual human cognition. Although this research focused on a specific byproduct of conceptual processing (i.e., NVS) and its accompanying neural components, we recognize the complex and dynamic processing within the brain resulting in the human ability to operate on abstract concepts.

In our results we seemingly uncovered an interplay between the type of memory search (open vs. structured) and the instruction to maintain fixation, which cannot be fully explained based on our data. Restricting NVS did not substantially impair memory performance in the open questions task, suggesting they are a byproduct of conceptual processing rather than a function. Yet, restricted gaze resulted in worse performance in structured retrieval task, meaning that either gaze fixation may require additional cognitive demands, or NVS help in some types of conceptual processing. This remains to be determined.

Lastly, our control task results were clearly biased towards the visual saccade task and did not allow us to fully assess the role of working memory processing and maintenance in modulating prefrontal activity preceding NVS. We attempted to do it using the memory saccade task, which was strongly underpowered in our case. Despite this, the data we obtained suggest that brain activity preceding NVS is qualitatively different from that in the memory saccade task and does not merely represent maintenance of mnemonic content. Further research is required to assess this complex processing network, and we believe NVS can be a good behavioral model for that. The interplay between NVS and content reproduction type also requires further investigation.

## Conclusions

In recent years, multiple authors have suggested that conceptual knowledge builds on sensorimotor circuitry in the brain, operating in low-dimensional conceptual spaces organized similarly to the representation of physical space. The available evidence suggests that organizing, storing, and manipulating conceptual knowledge is shared by temporal and parietal systems, with the involvement of other areas, such as the frontal cortex, being less well understood. Through the use of non-visual saccades as a behavioural model, our study extends established views on the organization of human conceptual knowledge by showing that the premotor cortex is involved in exploring and manipulating items within conceptual space.

## Acknowledgments

The authors would like to acknowledge support from Bial Foundation (grant 260/22) and Portuguese Foundation for Science and Technology (2023.08332.CEECIND) to AP, and Deutsche Forschungsgemeinschaft grant (122679504—SFB 874) to CK. We want to thank Alexandra Sipatchin and Fernanda Ponce for their help with eye tracking data analysis, Diogo Branco for his help with the task and the setup, Soraia Oliveira for help with data collection and Marcin Leszczynski, Susanne Dyck, Jon Walbrin, Fredrik Bergstroem, Elizabetta Del Re and Pieter Medendorp for helpful input and discussions.

## Author Contributions

AP conceptualized the experiment. AP, MJM, EA and GB designed the experiment and collected the data. LW, GB, AP and MJM analyzed the data. AP, LW, GB and MJM wrote the manuscript. All authors revised the manuscript.

## Data availability statement

Data underlying all figures is available for download at: https://osf.io/8q4wh/?view_only=730dcb4c2cab415694c3f7989241f31a

## List of Supplementary Materials

1. Supplementary Methods
2. Supplementary Results

