## Supplementary Materials for "Spontaneous saccades reveal that human premotor cortex is involved in conceptual processing"

#### 1. Supplementary Methods

Table S1. 40 Questions Stimuli

Block 1

- In which country were the Olympic Games born? [Information]
- What are the similarities between an orange and a banana? [Similarities]
- At which temperature does water boil in normal conditions? [Information]
- What are the similarities between a fork and a spoon? [Similarities]
- In which direction does the sun set? [Information]
- What is a thermometer? [Information]
- What are the similarities between a dog and a lion? [Similarities]
- In which continent is the Sahara desert? [Information]
- What are the similarities between yellow and green? [Similarities]
- What are the similarities between a coat and a shirt? [Similarities]

Block 2

- Refer to one of the constitutive powers of a democracy. [Comprehension]
- What is the main theme of the book of Genesis? [Information]
- What is the capital of Sweden? [Information]
- What are the similarities between the eye and the ear? [Similarities]
- What are the similarities between joy and sadness? [Similarities]
- What are the similarities between a car and a boat? [Similarities]
- What are the similarities between a piano and a drum? [Similarities]
- To which scientist is the “theory of relativity” associated? [Information]
- Who was Mahatma Gandhi? [Information]
- What are the similarities between a table and a chair? [Similarities]

Block 3

• Refer to some reasons that justify the importance of the study of History. [Comprehension]

• For what reason do deaf people have difficulty in learning to speak? [Comprehension]

• What are the similarities between Democracy and Dictatorship? [Similarities]

• What are the similarities between a fly and a tree? [Similarities]

• What are the similarities between an egg and a seed? [Similarities]

• What are the similarities between a poem and a statue? [Similarities]

• Why are defendants considered innocent until proven guilty? [Comprehension]

• Why do we pay taxes? [Comprehension]

• What are the similarities between fog and steam? [Similarities]

• Why is land more expensive in the city than in the countryside? [Comprehension]

Block 4

• What are the similarities between work and game? [Similarities]

• What are the similarities between praise and punishment? [Similarities]

• Why is the freedom of press important in a democracy? [Comprehension]

• Why do we need medical prescriptions to buy remedies? [Comprehension]

• Why do we wash our clothes? [Comprehension]

• What are the similarities between a friend and a foe? [Similarities]

• Refer to some reasons why we need to cook some foods. [Comprehension]

•For what reason do some people prefer to rob a bank rather than a friend? [Comprehension]

• What are the similarities between a coat and a suit? [Similarities]

• What are the similarities between hibernation and migration? [Similarities]

*Note: The above questions were originally in Portuguese as found in the Portuguese adaptation of WAIS-3 scale.*

**Eye tracking recording and preprocessing**

Eye movements were recorded at 100 Hz using the HTC VIVE Pro Eye (HTC Corporation, Taoyuan City, Taiwan) which features an eye tracker embedded in a virtual-reality-based head-mounted display. VR environments, which served as the stimuli described earlier, were developed, and an eye tracking software algorithm was implemented in Unity, using the SRanipal software development kit (SDK) provided by the HTC Corporation. For both eyes, pupil diameter, a measure for the validity of eye data, and the normalized eye gaze vector based on a right-handed coordinate system were extracted and averaged over both eyes. This method proved more robust than relying on a single eye metric. In this coordinate system, positive x- and y-values indicate a gaze towards the top-left, while positive z-values indicate a forward direction.

MATLAB (The MathWorks Inc., 2023) was used for preprocessing the eye data. After importing the raw eye data, the stability of the sampling rate was checked, and data and events were cleaned. Data was considered invalid for eye data validity values below 31. Various lowpass filter settings were tested. Due to the effect of Gibb’s phenomenon on square wave shapes such as saccades, lowpass filtering often introduces spikes at the edges of the signals.

Even with additional filters, spike residues were still identified. A Savitzky-Golay filter, as suggested by Nyström and Holmqvist (Nyström & Holmqvist, 2010), was tested and did not introduce spikes, but a better fit regarding the main sequence was still observed with a median filter due to a drastic decrease of velocities as a consequence of the Savitzky-Golay filter (Mack et al., 2017; Stuart et al., 2019). Ultimately, a median filter of length 10 was used to effectively remove spike noise while preserving the signal shape and saccade onset and offset points (Imaoka et al., 2020). This method also worked best on noisy data, reducing type I errors by accurately detecting saccade velocities without mistaking noise for saccades. Furthermore, amplitude calculations were most precise as onsets and offsets no longer fluctuated or contained overshoots at signal edges. Finally, a rather low velocity threshold could be applied, detecting small saccades while minimizing type I errors due to strong median filtering.

Saccades were detected using a velocity-based algorithm with a threshold of 30 deg/s (McPeek et al., 2000).

Since the recorded eye gaze vectors were normalized between -1 and 1, the angle between vectors was calculated as:

$$\alpha_{A,B} = \frac{180}{\pi} \cdot tan^{-1}\left(\frac{|A \times B|}{A \cdot B}\right),$$

with A and B containing two subsequent vectors with eye data positions x, y, and z. Velocity was calculated using the angle multiplied by the sampling rate. Saccade amplitude equaled the absolute value of α between saccade onset and offset.

During blinks, the pupil cannot be detected by the eye tracker, setting eye data validity values to 31. Blinks were detected using data marked as invalid. Rapid increases in vertical velocity due to Bell's phenomenon were observed before and after blinks. To avoid those artifacts being detected as saccades, data 100 ms before and after blinks was removed. Blink duration thresholds were set to 80 and 1000 ms (Hollander & Huette, 2022). Invalid data exceeding this threshold was considered missing. Data 100 ms before and after missing data was removed as well to account for any artifacts (Nij Bijvank et al., 2018). Invalid data for durations less than 80 ms was interpolated using spline interpolation, including single missing data points, to avoid rejecting saccades due to invalid data points. Trials with more than 30 % missing data, excluding blinks, were excluded from analyses. For the main task, each trial included markers for the beginning of the trial, the end of the question, and the end of the trial, including information about whether the answer was known. Only trials with a full set of events were included.

Saccade onsets were marked before an average of two subsequent samples exceeded the specified threshold. Given the variability of saccade slopes, for a more precise onset, the last local minimum below 10 deg/s in an interval of -2 to +1 data points relative to the previously marked onset was used as the final saccade onset. For exported saccade events to EEG data, more precise saccade onsets were calculated by interpolating the velocity data to 500 Hz. Onsets were then calculated using a procedure analogous to the one described above, with adjustments made to the intervals to match the new sampling rate. Saccade offsets were marked after the average of two subsequent samples fell below the threshold. Saccade duration thresholds were set to 18 to 240 ms, saccades above or below the threshold were removed. Due to a minimum refraction time of 20 ms following saccades, saccades with inter-saccadic intervals of less than 20 ms were combined (Larsson et al., 2013; Nij Bijvank et al., 2018). Additionally, saccades containing invalid data were excluded. The main sequence, representing the relationship between saccade amplitude and peak velocity, was also used as a filtering criterion. A second-degree polynomial fit was applied to the main sequence for each participant. Outliers exceeding 4 SD from the fit were detected and removed, as they most likely do not represent saccades, but noise. For the EEG analysis, epochs selected were only those presaccadic periods that were not preceded by saccades or blinks within 200 ms. This naturally controlled for eye movement artifacts, eliminating the need for ICA during EEG processing, which would have in turn risked removing relevant data, while no increase in the SNR of the studied contrasts has actually been found when specifically tested (Delorme, 2023).

Fixations were extracted using a velocity threshold below 20 deg/s and a minimum duration of 200 ms (Liu et al., 2018). Data points within 150 ms at the beginning and end of fixations were removed. Since accuracy is not of interest in this particular case, precision metrics, including RMS and SD, were calculated, and outliers were used as exclusion indicators. Main sequences were plotted for each participant followed by careful visual inspection, and abnormal distributions or lack of identifiable saccadic profiles, indicating noise, were grounds for exclusion. Participants with fewer than 30 valid trials were excluded. Twelve participants were excluded from the behavioral and EEG analysis. The final sample size included 45 participants, with 24 in the fixed condition and 21 in the free condition.

For the control task, the velocity threshold for saccade detection was maintained at 30 deg/s to accurately detect and remove all saccades preceded by eye movement within 200 ms and control for those artifacts in the EEG analysis. Since low saccades would not reach the predefined target, saccades below 150 deg/s were excluded as well. Execution results were classified as correct or incorrect based solely on the x-coordinate. An event marker within the data indicated cue to start executing a saccade when the fixation cross disappeared. If, within the interval of -500 to 1250 ms, at least 20 ms were consistently, depending on the condition, above or below a threshold of 0.13 relative to the average x-baseline, the trial was marked as correct. Due to the removal of saccades preceded by other eye movements, a trial can still be rejected if it is executed wrongly, but directly afterwards executed correctly.

The saccade slope was calculated to evaluate if the saccade was executed in the correct direction. Only saccades in the correct direction and above or below the baseline-corrected x-value were included. This ensured that saccades moving from the wrong target back to the center were excluded, although indicating a correct slope. Overall, only saccades toward the targets and not back to the center were included. For the visual saccade task all saccades from the blink signal to the end of saccade cue interval were included, accounting for system delays by including 100 ms prior.

The task was initially designed to focus on saccades following the saccade cue, aiming to avoid overlapping visual responses and motor programming. However, due to inadequate instructions, participants executed saccades before the saccade cue. Although analyses revealed no statistical difference between saccades executed prior to and after the saccade cue, we utilized the memory task to more specifically differentiate between visual responses and motor programming. For the memory task, only saccades occurring 100 ms after the saccade cue were included, ensuring the task remained a memory task rather than a visual task with visible targets. Trials 50 and 100 were excluded for all participants due to a technical error. Invalid trials, characterized by no detected saccade, saccades in the wrong direction, or more than 30 % missing data, were also excluded. Overall, 3 participants were excluded from the EEG analysis for the control task due to these criteria. For the memory task, participants with fewer than 25trials were excluded from the analysis. Due to inadequate instructions, a larger number of participants had to be excluded from analyses involving the memory task. The final sample sizes were determined after EEG processing.

**EEG recording and preprocessing**

EEG data was recorded using either a 64-electrode dry EEG or a 256-electrode wet EEG system (ANT Neuro, Enschede, Netherlands). Initially, the 256-channel system was employed for data collection. However, due to unavoidable technical difficulties encountered during the recording process, the remaining participants were recorded using the 64-electrode system to ensure the continuation of the study without compromising data collection. Electrodes were arranged in an equidistant montage using an elastic cap. For the 64-channel setup, electrodes on the left and right mastoids acted as online ground and reference, respectively. In the 256-channel setup, CPz and an electrode on the left mastoid were used as online reference and ground. Impedances were kept below 25 kΩ. The signals were recorded with a sampling rate of 500 Hz.

Data preprocessing was performed using EEGLAB (Delorme & Makeig, 2004) and MNE-Python (Gramfort, 2013; Larson et al., 2024). After importing the raw EEG files and the channel locations, the stability of the sampling rate was checked. A high-pass filter of 1 Hz and a low-pass filter of 40 Hz were applied (Kovalenko & Busch, 2016). After visual inspection, environmental noise became evident at 25 Hz which was effectively removed using a notch filter of 25 ± 0.2 Hz. A total of 11 out of 29 participants were affected by this noise to varying extents, with all channels impacted in 5 participants, approximately 50 % of channels affected in 2, and around 10 % of channels affected in 4 participants. All filters used were zero-phase hamming windowed sinc FIR filters. Passband edges are specified instead of cutoff frequencies with a transition band width of 25 % of the lower passband edge, but not lower than 2 Hz. Due to soft roll-off, remaining line noise was additionally removed prior to low-pass filtering using a notch filter of 50 ± 2 Hz. Flatline channels were removed using EEGLAB's pop_clean_rawdata function if a channel was flat for more than 5 s (Delorme, 2023). Additionally, noisy channels were rejected, using EEGLAB's pop_rejchan function based on kurtosis (4σ), probability (4σ), and abnormal spectra (1 – 40 Hz, 3σ), followed by manual rejection based on careful visual inspection (Brandmeyer & Delorme, 2020; Rodrigues et al., 2021). Removed channels were interpolated using spherical spline interpolation. Finally, the data was re-referenced to the average of all electrodes. ICA was not applied to avoid possible removal of saccade-planning data. Instead, it was controlled for eye movements prior to intervals of interest using eye tracking data.

EEG events were cleaned and processed separately for the main and control task. For the main task, saccade events were imported from eye tracking data, using trial onsets for synchronization. For the main task, alignment offsets were kept below 8 ms for all trials with a mean offset of 2.6 ± 0.5 ms. Trial markers included trial onset, end of question, and beginning of the verbal answer period. Trials lacking any essential markers are excluded. Additionally, only saccades occurring after the end of the question and before the verbal answer period were analyzed, all other saccades were excluded. This approach aimed to control for the impact of auditory and language processing as well as speech-related artifacts on the EEG data. For the control task, each of the 4 blocks was synchronized separately, due to breaks during recording. For the control task, event markers were not stored separately, but within the eye data and are therefore limited to the sampling rate of the eye data. Therefore, occasionally, single trials exhibited alignment offsets larger than 10 ms, up to 16 ms. Since this affected approximately only 2 % of trials on average, the impact was deemed negligible. Otherwise, alignment offsets were kept below 10 ms for all trials, with a mean offset of 4.0 ± 0.4 ms. Since saccades for the control task had already been extensively processed within the eye data processing, no additional saccades were excluded.

Epochs were extracted from the continuous EEG data using a time interval from -200 to 100 s relative to the onset of a saccade. During the import of saccade events from processed eye data, only saccades that were not preceded by saccades or blinks within 200 ms prior to saccade onset were considered to control for eye movement artifacts. The interval from -200 to -170 ms was used for single-trial baseline correction (Meghanathan et al., 2020; Nikolaev et al., 2013; Revankar et al., 2020). The baseline correction was performed using a single trial normalization instead of the trial average. Epochs were rejected based on the interval -200 to -20 ms prior to saccade onset to exclude spikes originating from oculomotor activity (Revankar et al., 2020). Semi-automatic artifact rejection occurred if the amplitude exceeded 50 µV between two subsequent samples or if the amplitude difference exceeded 100 µV in an interval of 50 ms (Meghanathan et al., 2020; Nikolaev et al.). Four regions of interest (ROIs) were defined: frontal left (FL), frontal right (FR), parietal left (PL), and parietal right (PR), distributed around corresponding electrodes F3, F4, P3 and P4 in the 10–20 electrode system, as seen in Figure S1.

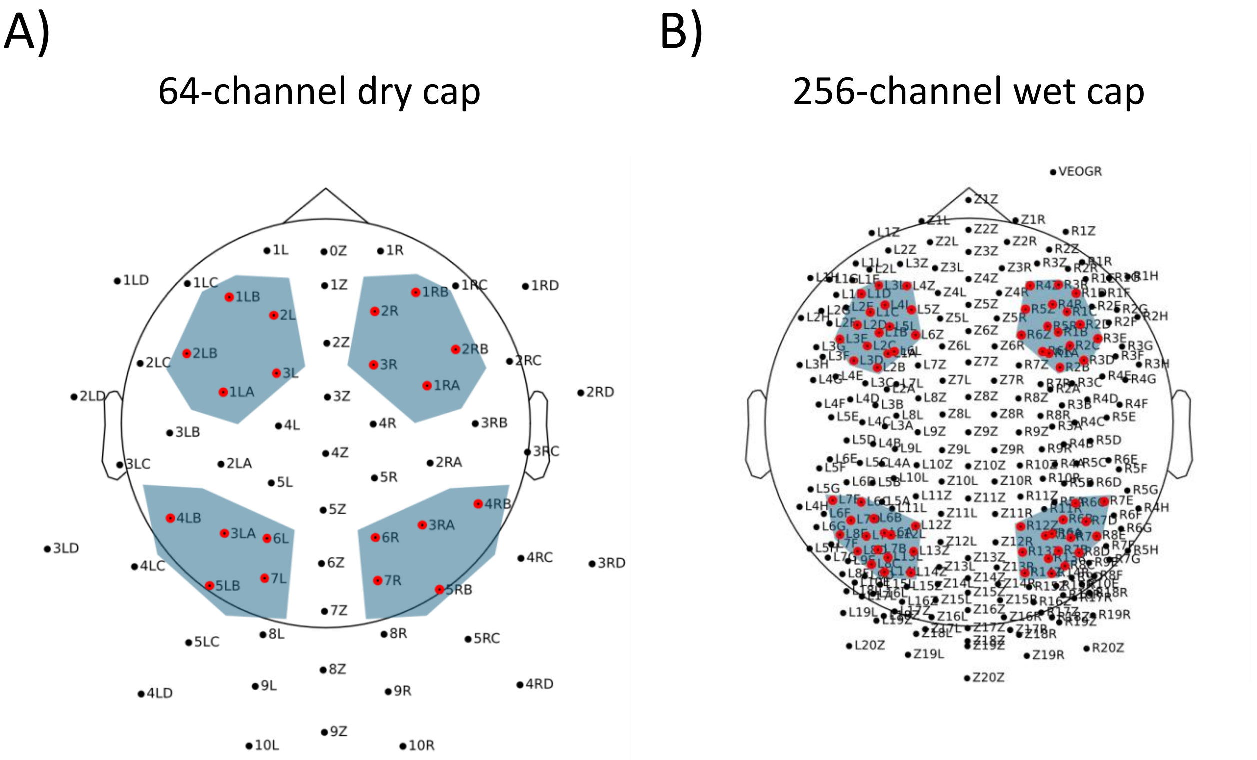


Figure S1. Detailed electrode layout highlighting ROIs (FL, FR, PL, PR) in blue. A) 20 electrodes ROI for 64-channel system. B) 64 electrodes ROI for the 256-channel EEG systems.

Since the FEF and PPC are the primary areas of interest in saccade research and are both superficial and spatially distinct, this setup allowed us to effectively distinguish neural activity in these regions. Our analysis focused on these areas due to their critical roles in saccadic control and cognitive processing during the tasks. Each ROI included 5 electrodes for the 64-channel and 16 electrodes for the 256-channel EEG system. To ensure consistency between the 64-channel and 256-channel EEG systems, ROIs were matched using 3D scalp topographies. This involved normalizing channel locations across both systems, overlaying them, and identifying corresponding ROIs based on the electrode positions. For ERP visualization, electrodes from the 64-channel system were interpolated to match the 256-channel system's locations using only the preselected ROI and EEGLAB’s integrated spherical spline interpolation (Freeden, 1984; Kang et al., 2015; Perrin et al., 1989), a method that has been applied before for upsampling due to its ability to handle any electrode arrangement while maintaining data integrity (Abdoun, 2011; Nouira et al., 2016). This method was not intended to provide new information but to make the data comparable to the 256-channel system for accurate ERP visualization and comparison between cap configurations, which is why more advanced methods, like Convolutional Neural Networks (CNNs) for upsampling, commonly used today to increase information (Svantesson et al., 2021), were not needed and therefore not employed. For statistical analyses, the original data was used.

Only participants with complete datasets, including eye data for the main and control tasks and EEG data for both, were included in the analysis. Several participants had to be excluded due to incomplete EEG data, such as the loss of triggers from the eye tracker computer to the EEG. Participants with fewer than 40 trials for each main and control saccade condition were also excluded. Due to prior trial rejection, this exclusion criterion effectively removed participants with noisy data. Overall, the final sample size included 29 participants (mean age 26.9 ± 7.0 years, 11 male, none left-handed). The final trial count for main tasks was 2884 (mean 99.5 ± 52.26 per participant) and for control visual tasks was 1777 (mean 61.28 ± 16.79). The final sample included 11 participants recorded using a wet cap, 18 using a dry cap, 17 in the fixed condition, and 12 in the free condition. For analyses involving the memory task, the same procedure and similar exclusion criteria were applied. However, due to inadequate task instructions, more participants had to be excluded. Participants with more than 25 trials were included, resulting in a final sample size of 12 participants (mean age 25.3 ± 5.7 years, 4 male, none left-handed). The final trial count for control memory tasks was 391 (mean 32.58.5 ± 7.11 per participant).

Voxel-based volume source reconstruction was performed on the main, visual, and memory saccade tasks to evaluate the source activity observed in the ERP data using a template MRI. Using the interpolated 256-electrode data was not feasible for source reconstruction, as interpolations introduced errors in the covariance matrix computations, leading to unreliable results. To address this issue, the 256-electrode system was downsampled to match the 64-electrode layout. The three-layer boundary element model (BEM) in the MNE-Python toolbox, along with its corresponding volume source space, was used for source reconstruction. Digitized electrode locations were aligned to the generic head model to compute a forward solution based on the template MRI. Noise covariance matrices were computed for each participant and condition based on the presaccadic baseline interval -200 ms to -170 ms relative to saccade onset of the corresponding epochs. Inverse operators were created for each participant and condition using the forward solution and noise covariance matrices. The forward model and the ERP for each condition were then used to estimate an inverse model using the standardized low-resolution brain electromagnetic tomography (sLORETA) method (Pascual-Marqui, 2002). Source estimates were subsequently averaged over a specified time window 150 ms to -20 ms relative to saccade onset (Abe et al., 2004).

**Behavioral analysis**

For the behavioral analysis of the eye data recorded during the main task, answer duration, saccade and blink frequencies, pupil diameter, saccade amplitudes, peak velocities, and angles were extracted for each trial and participant, focusing on the period after each question until the end of each trial. Since saccade amplitudes and peak velocities are closely correlated according to the main sequence, only saccade amplitude was analyzed.

Saccade angles were classified and analyzed relying on their absolute offset-coordinates. Saccades were divided into four quadrants using a coordinate system rotated by 45 degrees, with the first quadrant being right, the second top, the third left, and the fourth down (see Figure S11 B). This approach allowed for the analysis of saccade distribution within the field of view for behavioral analyses and highlighted potential biases within the EEG data. Slight variations in main clusters around the origin indicated eye tracker calibration offsets. To address potential misclassification of saccades, a hierarchical clustering algorithm identified the central cluster around the data median, and its centroid was calculated separately for each participant. Saccades were centered based on the centroids. Subsequently, saccades near the center were excluded by removing those within ± 1.5 SD from the center for both x- and y-directions. Saccade counts for each factor combination were normalized using total saccade counts. Angles were regarded from the observer's point of view. For the statistical analysis, direction as an additional fixed factor was included in the model. This factor comprised four instead of two levels, increasing the model complexity.

Parameters were averaged across combinations of factors for each participant, specifically blocks 1-4, condition type (fixed/free), and question type (open/structured). Due to a technical error, the "don't know" responses included both incorrect answers and questions answered with "don't know." Incorrect answers likely involve a different level of cognitive engagement compared to simple "don't know" responses. When participants provide incorrect answers, it suggests that they are attempting to retrieve or deduce the correct information, engaging their cognitive processes even if the final response is wrong. In contrast, "don't know" responses can occur for various reasons, including a lack of effort or genuine uncertainty. Since our interest lies in cognitive engagement, including "don't know" responses could confound the results due to the mix of engagement during responses. Due to the lack of consistency regarding the classification of "don't know" answers, only known answers were included in the main analyses. Separate analyses assessed the "answer known" factor to avoid model complexity from including too many factors. Additionally, the percentage of "don't know" responses varied across trials and factors, adding inconsistency. This non-linear distribution of "don't know" responses further justified their exclusion to maintain the integrity of assessing cognitive engagement.

JASP (JASP Team, 2024) was used for statistical analyses. Extreme values of averaged parameters were identified as outliers and excluded using the interquartile range (IQR) method, with a multiplier of 1.5 times the IQR. Normality tests, specifically the Shapiro-Wilk and Kolmogorov-Smirnov tests, were applied. The Shapiro-Wilk test revealed non-Gaussian distribution for all parameters, while the less conservative Kolmogorov-Smirnov test indicated normal distribution for saccade frequency and pupil diameter. To account for varying trial ratios and include random effects, a generalized linear mixed model (GLMM) was chosen as an alternative to repeated measures analysis of variance (ANOVA) (Casals et al., 2014). Since GLMMs are reported to be robust to non-normally distributed data and marginal distributions are only one of many indicators for model selection, analyses using a gamma distribution with a log link function and a Gaussian distribution with an identity link function were compared. Since both displayed similar goodness-of-fit parameters (AIC, BIC), further, corrected Pearson residuals of Gaussian GLMM were assessed for normality using Q-Q plots and subsequently statistically tested for normality as indication of model fit (Agresti, 2007; Anderson et al., 2012; Cordeiro & Simas, 2009; Dunn & Smyth, 2018; Schielzeth et al., 2020; Salinas et al., 2023; Xia & Sun, 2023). Q-Q plots showed constant variance. A Kolmogorov-Smirnov test confirmed normality of residuals parameters, indicating homoscedasticity for all parameters (Schielzeth et al., 2020). Given GLMM's robustness and indicators of a sufficiently good model fit, ultimately, a Gaussian GLMM with an identity link function using the likelihood ratio test was applied.

Due to non-normal marginal distributions, the Wilcoxon signed-rank test was used for post hoc comparisons as a non-parametric alternative to paired sample t-tests. For condition type as between-subject effect, post hoc tests using the Wilcoxon rank-sum test were employed due to uneven and unpaired datasets, serving as an alternative to independent t-tests. Holm correction was used to account for multiple testing. Additionally, since condition type did not vary across the random effects, the GLMM often did not pick up on significant differences, but post hoc tests did. Generally, when including random effects for participants, the model accounts for the variability between participants. If a fixed effect such as groups between participants does not vary within participants, its significance can be overshadowed by the variability attributed to the random effects. To account for this effect, post hoc tests were taken into account for condition type even if GLMM was not significant.

**EEG analysis**

Statistical analysis of ERP data was additionally performed on separate ROIs. Averaged EEG amplitudes of conditions were statistically compared using 4 × 2 repeated measures ANOVAs with 4 ROIs and condition type as factors. For analyses of between-subject conditions such as fixed versus free conditions or wet versus dry cap, ANOVA was used to account for trial imbalances. Post hoc independent t-tests were applied with Holm correction for multiple testing. To assess the statistical significance of the source activity, source estimates for each condition were extracted. Spatial adjacency for the source space was computed to enable cluster-based permutation testing. A non-parametric, two-tailed cluster-based permutation test was performed on the data, involving voxel-wise comparisons to zero and inherently correcting for multiple testing (Abe et al., 2004). Statistical significance was assessed using a p-value threshold of 0.05. T-values were projected into sLORETA images to visualize the source activity, highlighting regions with significant neural dynamics. Anatomical labels were obtained using the FreeSurfer aparc parcellation scheme (Desikan et al., 2006) to map source activity to specific cortical regions.

It should be noted that only cortical labels will be returned. The focus on cortical areas aligns with the goal of analyzing cortical involvement in saccadic movements, particularly in distinguishing activity between visual and non-visual saccades. Subcortical areas, while important, were not directly captured due to the limitations of the parcellation used. Source estimates were aligned with these labels to investigate neural activity in distinct areas. For each significant cluster identified in the permutation test, the t-values of voxels were examined relative to the anatomical labels, and the maximum t-value for each label was extracted.

### 2. Supplementary Results

#### Saccade Metrics

The plots in Figure S2 (A and B) show the distributions of non-visual saccade characteristics during the main saccade task, based on all saccades exported to the EEG pipeline (n=29). All figures show only the most relevant ranges of data after processing, with only detected saccades above 30 deg/s included. However, it is important to note that while the plots in Figure S2 reflect all detected saccades during the full answer period after participants were asked a question, the EEG analysis focused on a narrower time window - specifically, the presaccadic interval before participants began speaking. EEG preprocessing steps (i.e. exclusion of trials overlapping with speech onset or containing artifacts) were applied later, resulting in a smaller subset of trials used for final EEG analyses. Therefore, the histograms shown here represent the broader set of exported saccades and do not directly reflect the final trial counts included in EEG statistical analyses. The distribution of saccade peak velocity shows few saccades reaching up to 900°/s, with the majority of saccades below 600°/s. The distribution of saccade amplitude shows few saccades reaching up to 50° but most below 20°. The saccade angle distribution resembles a bimodal distribution, with clear peaks at cardinal and oblique directions and the majority of saccades directed along horizontal or near-horizontal axes.

The average distribution of saccade counts within each velocity-based bin as taken for EEG analyses is shown in Figure S2 C, indicating most saccades considered were above 100 deg/s.

The plots in Figure S2 (D and E) show the saccade metrics during the control task, based on all saccades exported to the EEG pipeline (n=29). While there was no need to restrict to a prespeech window as in the main task, the same EEG preprocessing steps (i.e. artifact rejection) were applied, meaning that the plotted distributions also reflect a broader exported set and not the final number of EEG trials analyzed. Since only saccades above 150 deg/s were extracted, the metrics are skewed towards larger saccades as seen in the peak velocity and amplitude. Saccade angles are directed along the horizontal axis only due to the task design specifically including saccades to the left and right side.

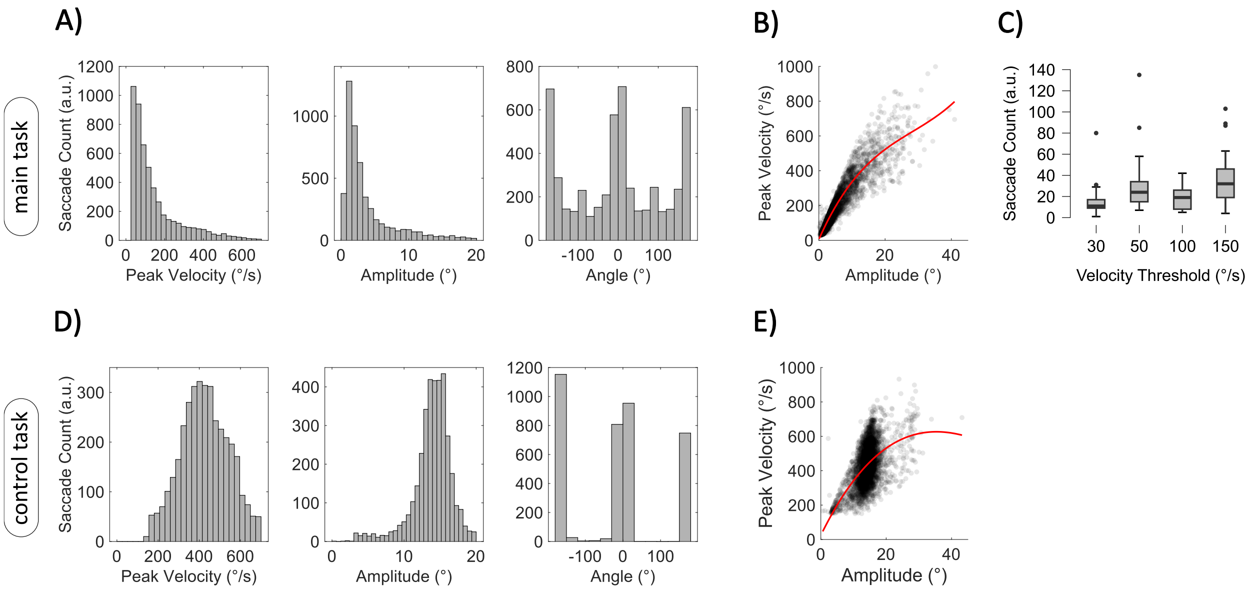


Figure S2. Saccade metrics during the main and control task prior EEG processing. A) Histograms of saccade peak velocity, amplitude, and angle across all saccades of the main task exported for the EEG analysis (n=29). B) Main sequence for all saccades of the main task exported for the EEG analysis (n=29), showing the relationship between saccade peak velocity and amplitude with a third-degree polynomial fit (red line). C) Average saccade counts across participants for the main task after EEG processing (n=29), shown at different saccade velocity thresholds. D) Histograms of control saccade metrics (peak velocity, amplitude, and angle) exported for the EEG analysis (n=29). E) Main sequence for control saccades exported for the EEG analysis (n=29).

**Comparison of wet and dry caps**

To compare ERPs across different tasks, a comparison between dry and wet cap EEG systems was conducted to reliably concatenate results for both caps. The effect of the EEG system on presaccadic potentials was assessed for the main and control tasks separately, using ANOVA with cap (wet/dry cap) and frontal/parietal ROI as factors (see Figure S3 A, B and Table S2, S3). For the main task, no significant main effect for condition was found ($F(1,54) = 0.958, p = 0.332, \omega^2 = 0.000$), nor was there a significant interaction effect for condition × ROI ($F(1,54) = 0.843, p = 0.363, \omega^2 = 0.000$). Similarly, for the control task, there was no significant main effect for cap ($F(1,54) = 5.681 \cdot 10^{-5}, p = 0.994, \omega^2 = 0.000$) and no significant interaction effect for cap × ROI ($F(1,54) = 0.850, p = 0.361, \omega^2 = 0.000$). Bayesian analysis further supported these findings, with $BF_{10} = 0.350$ for the main task and $BF_{10} = 0.273$ for the control task, indicating moderate evidence in favor of the null hypothesis and suggesting no effect of EEG cap. The analysis revealed that the type of EEG cap did not significantly affect presaccadic potentials in either the main or control tasks. No significant differences were observed in the main effects or interaction effects related to the EEG system. Therefore, analyses across different EEG cap types can be reliably concatenated.

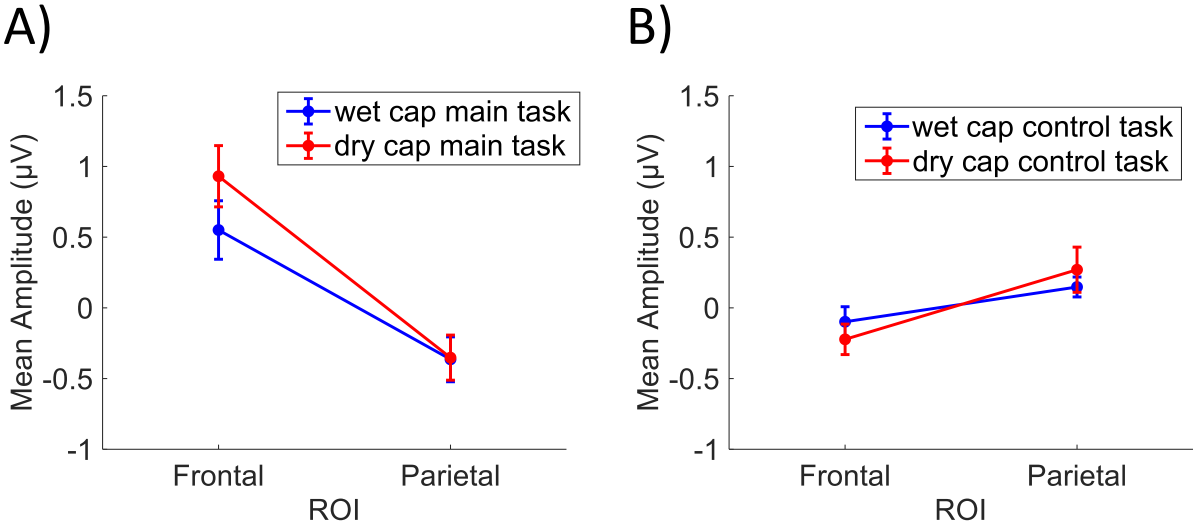


Figure S3. EEG potentials time-locked to saccades for preliminary analyses. Mean amplitudes averaged from  150 ms to  20 ms are shown across frontal and parietal ROIs with error bars indicating the standard error of mean. Mean amplitudes are shown for A) main task recorded with the wet and dry cap (n=11/n=18), B) control task recorded with the wet and dry cap (n=11/n=18).

Table S2. Summary of ANOVA for the effects of EEG-cap-condition (dry vs. wet) and combined ROI on averaged presaccadic EEG amplitudes during the main task (n=11/n=18).

| **Cases** | **Sum of Squares** | **df** | **Mean Square** | **F** | **p** | **ω²** |
|---|---|---|---|---|---|---|
| Condition | 0.525 | 1 | 0.525 | 0.958 | 0.332 | 0.000 |
| ROI | 16.479 | 1 | 16.479 | 30.049 | < .001 | 0.318 |
| Condition ✻ ROI | 0.462 | 1 | 0.462 | 0.843 | 0.363 | 0.000 |
| Residuals | 29.614 | 54 | 0.548 | | | |

Note. Type III Sum of Squares

Table S3. Summary of ANOVA for the effects of EEG-cap-condition (dry vs. wet) and combined ROI on averaged presaccadic EEG amplitudes during the control task (n=11/n=18).

| **Cases** | **Sum of Squares** | **df** | **Mean Square** | **F** | **p** | **ω²** |
|---|---|---|---|---|---|---|
| Condition | $1.385 \times 10^{-5}$ | 1 | $1.385 \times 10^{-5}$ | $5.681 \times 10^{-5}$ | 0.994 | 0.000 |
| ROI | 1.860 | 1 | 1.860 | 7.629 | 0.008 | 0.102 |
| Condition ✻ ROI | 0.207 | 1 | 0.207 | 0.850 | 0.361 | 0.000 |
| Residuals | 13.166 | 54 | 0.244 | | | |

Note. Type III Sum of Squares

**Preliminary EEG Analyses**

Following this, presaccadic potentials were analyzed for main versus control task, with control task including both visual and memory saccade tasks. Figure S4 A, B illustrates the grand average ERPs across all electrodes for both conditions. While this figure provides an overview of the data, it also demonstrates that the chosen ROI represents the largest task-related deviations, confirming that it effectively captures the key differences. Importantly, no other electrodes demonstrate more pronounced or relevant deviations, supporting the validity of our ROI selection in capturing the effects of interest.

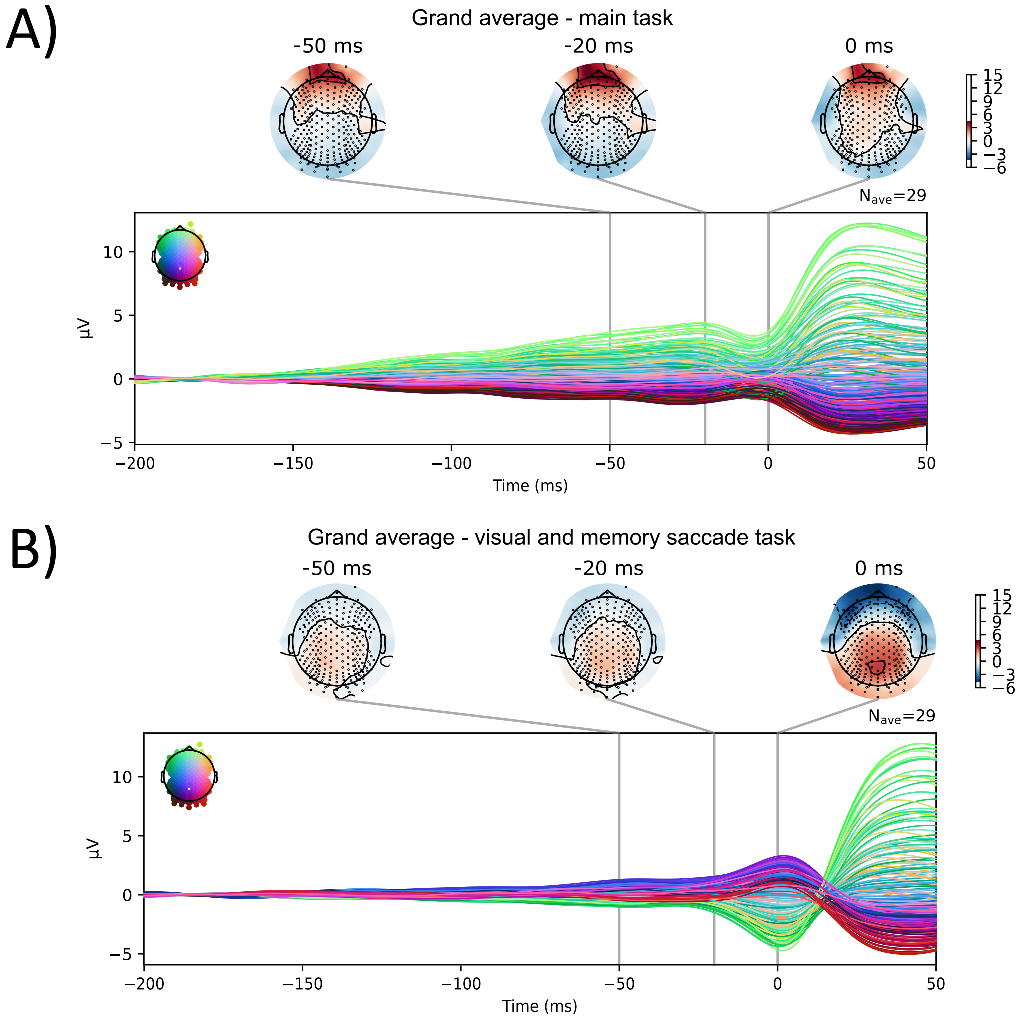


Figure S4. Grand average ERPs across all electrodes. Topographic plots at -50 ms, -20 ms, and 0 ms relative to saccade onset are shown (n=29). ERPs are shown for A) main task and B) VST and MST control tasks.

**Saccadic condition preselection**

For the control task, due to task conditions, only large saccades with a minimum velocity of 150 deg/s were included. In the main task, saccades of variable velocities and amplitudes were executed, with all saccades above 30 deg/s being included. Comparisons were conducted between EEG epochs time-locked to main task saccades below 50 deg/s and main task saccades above 150 deg/s using repeated measures ANOVA with velocity and ROI as factors to evaluate the effect of saccade amplitude on presaccadic potentials (see Figure S5 and Table S4). No significant main effect for velocity was found ($F(1,28) = 0.656, p = 0.425,\ \omega^2 = 0.000$), nor was there a significant interaction effect for velocity × ROI ($F(1,28) = 1.122, p = 0.298$, $\omega^2 = 0.002$). Bayesian analysis further supported these findings, with $BF_{10} = 0.201$ for frontal ROI and $BF_{10} = 0.919$ for the parietal ROI, indicating evidence in favor of the null hypothesis which further showed that saccade size did not have a significant effect on presaccadic potentials. Therefore, including saccades of varying amplitudes and velocities in the main task analysis - unlike the more uniform saccade sizes in the control task - can be justified.

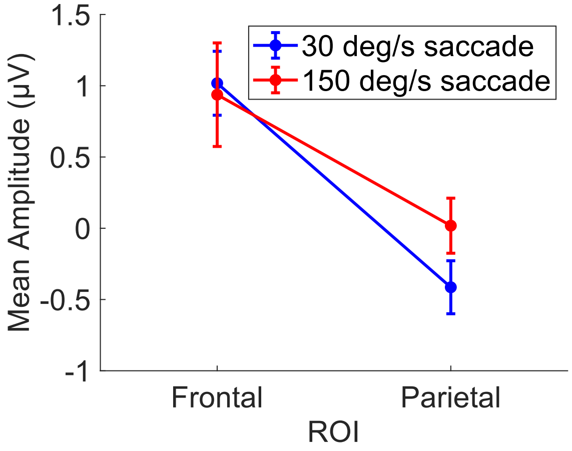


Figure S5. EEG potentials time-locked to saccades for saccadic condition preselection. Mean amplitudes averaged from  150 ms to  20 ms are shown across frontal and parietal ROIs with error bars indicating the standard error of mean. Mean amplitudes are shown for the main task with a minimum velocity of 30 deg/s and 150 deg/s (n=29).

Table S4. Summary of repeated measures ANOVA for the effects of velocity-condition (30 deg/s vs. 150 deg/s) and combined ROI on averaged presaccadic EEG amplitudes during the main task (n=29).

| Cases | Sum of Squares | df | Mean Square | F | p | ω² |
|---|---|---|---|---|---|---|
| condition | 0.897 | 1 | 0.897 | 0.656 | 0.425 | 0.000 |
| Residuals | 38.264 | 28 | 1.367 | | | |
| ROI | 40.059 | 1 | 40.059 | 19.951 | < .001 | 0.233 |
| Residuals | 56.221 | 28 | 2.008 | | | |
| condition ✻ ROI | 1.903 | 1 | 1.903 | 1.122 | 0.298 | 0.002 |
| Residuals | 47.466 | 28 | 1.695 | | | |

Note. Type III Sum of Squares

Additionally, main and control tasks were compared using repeated measures ANOVA with task and ROI as factor, but including only saccades above 150 deg/s to further justify the inclusion of all saccades sizes within the main task. Once again, a significant main effect of "task" ($F(1,28) = 5.031, p = 0.033, \omega^2 = 0.066$) was found. Additionally, a significant main effect of "ROI" ($F(1,28) = 7.538, p = 0.015, \omega^2 = 0.126$) was reported. A significant interaction effect of task × ROI ($F(1,28) = 24.631, p < 0.001, \omega^2 = 0.365$) could also be found. Analyses restricted to saccades above 150 deg/s yielded the same statistical results as those including all saccades, further justifying the inclusion of smaller saccades in our main task analysis compared to those in the control task. The results indicate that the ERP effects observed are not driven by differences in saccade velocity or amplitude between the main and control tasks.

**Additional Analyses of Main and Control Tasks**

Figure S6 A, B provides an extended view of the EEG results for the main and control tasks, covering a wider temporal range than the main analysis. No baseline correction was applied, allowing for a direct representation of EEG activity without imposing a predefined reference period. This approach underscores that all reported main findings represent relative activity changes rather than absolute activity levels, given their interpretation relative to a baseline.

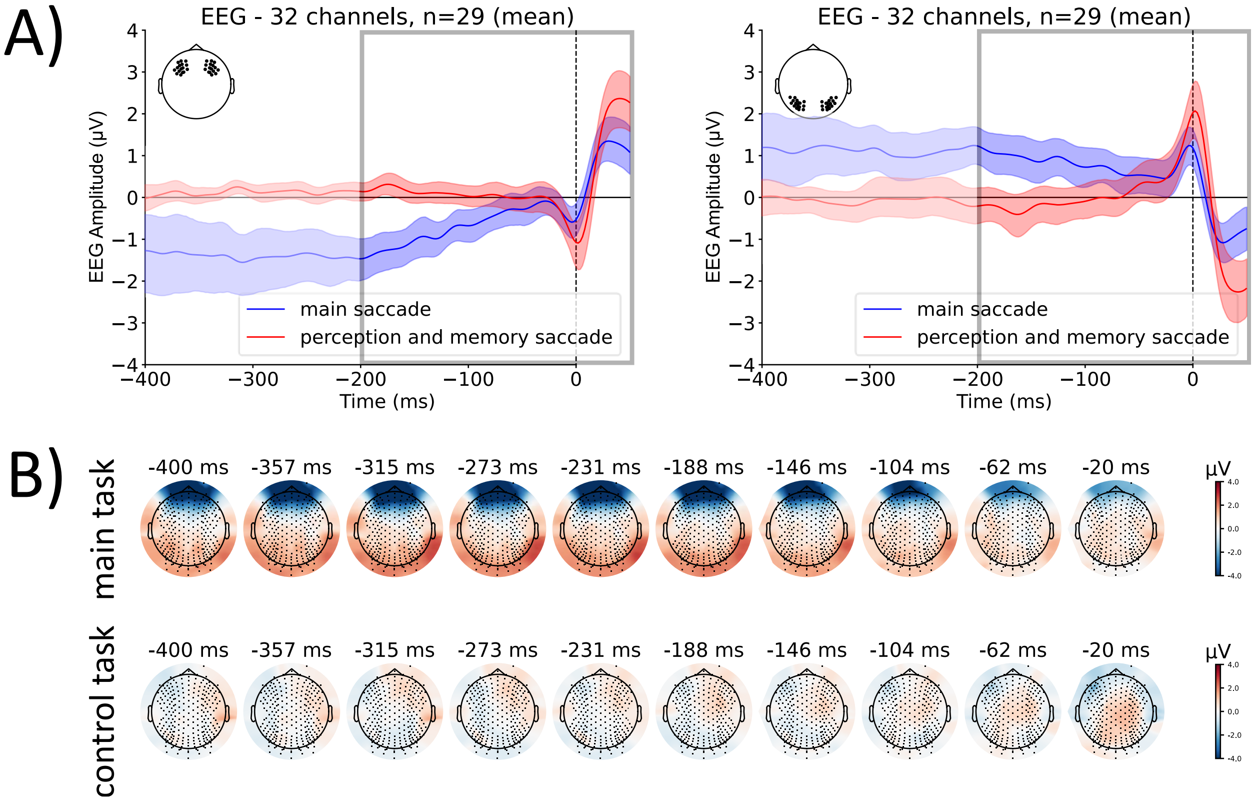


Figure S6. EEG results for main vs. control tasks over an extended time window without baseline correction. A) ERPs relative to saccade onset for main and the control tasks; shaded areas denote 95% confidence intervals; outlined grey box denotes the time window used in the main analysis; B) Topographic maps of presaccadic activity across main and control tasks.

In addition to the sliding-window analysis comparing ERP amplitudes between the main and control tasks, we also tested whether ERP amplitudes in each condition were significantly different from baseline. For this, a sliding-window analysis of one-sample t-tests against zero was performed. For the main task, significant deviations from baseline were observed in the frontal ROI at -150 ($p = 0.038$) and in the parietal ROI at -120 ms ($p = 0.028$). For the control task, significant deviations occurred in the frontal ROI at -70 ms ($p = 0.041$) and in the parietal ROI at -60 ms ($p = 0.029$). These results confirm that ERP deflections in both tasks were statistically distinguishable from baseline, complementing the condition-comparison analysis presented in the main manuscript.

To test whether the effects observed for frontal and parietal ROIs are lateralized across left and right channels, we conducted additional repeated measures ANOVA with task and ROI

(FL/FR/PL/PR) as factors. Once again, a significant interaction effect of task × ROI ($F(2.730,76.451) = 13.013, p < 0.001, \omega^2 = 0.272$) was reported. However, post hoc tests identified no significant differences between the left and right ROI for the main task's frontal ($p = 1.000$) and parietal ($p = 1.000$) regions, nor for the control task's frontal ($p = 1.000$) and parietal ($p = 1.000$) regions, confirming no lateralization of the effect seen prior.

Subsequently, visual and memory saccade control tasks were compared separately to the main task allowing for differentiations between visual and motor programming effects on presaccadic potentials. The grand average ERPs and topographic visualizations for the visual control task for all participants demonstrate similar patterns as described earlier with slightly less left-lateralization seen for parietal regions (see Figure S12 A-D). Comparing main and visual saccade task using combined frontal and parietal ROIs, a significant main effect of task ($F(1,28) = 17.059, p < 0.001$, $\omega^2 = 0.144$) and ROI ($F(1,28) = 5.735, p = 0.024, \omega^2 = 0.119$) was found. Additionally, a significant interaction effect of task × ROI ($F(1,28) = 25.871, p < 0.001, \omega^2 = 0.414$) was found. Post hoc tests showed identical results to the comparison between the main task and overall control task. Similarly, the repeated measures ANOVA analysis with separate ROIs for lateralization assessment yielded the same results. This outcome was expected since visual saccade tasks dominate the combined control tasks, resulting in a higher proportion of visual tasks compared to memory tasks in the control condition.

The ERP results for the main and memory control tasks for 12 participants demonstrate a high degree of similarity, with key differences still identifiable (see Figure S12 E-H). Specifically, the main task shows a larger positive potential over frontal regions, while more pronounced negative activity is seen over parietal areas compared to the visual control task. The average mean amplitudes suggest that frontal and parietal regions exhibit nearly identical potentials, which contrasts with the visual control task. The main task retains previous characteristics, though greater variability is observed, likely due to the smaller participant sample size. The topographic map for the control task highlights both frontal and parietal potentials, with the parietal potential appearing more widely distributed and the frontal potential being more localized. Additionally, a cluster is visible in the left hemisphere of the centro-frontal region, similar to what was observed in the visual and combined visual-memory saccade control tasks. However, new bilateral frontal clusters are also noted, indicating distinct neural patterns associated with the memory control task. As before, no dominant lateralization can be observed. For the comparison between main and memory task, only 12 participants were included. Repeated measures ANOVA with combined ROIs as factors revealed a significant main effect of task ($F(1,11) = 5.055, p = 0.046, \omega^2 = 0.090$), with medium to large effect size. No significant main effect was found for ROI ($F(1,11) = 3.361, p = 0.094, \omega^2 = 0.147$). There was also no significant interaction effect of task × ROI ($F(1,11) = 1.418, p = 0.259, \omega^2 = 0.031$). Repeated measures ANOVA with separate ROIs showed neither a significant main effect of ROI ($F(2.018,22.200) = 2.473, p = 0.107$, $\omega^2 = 0.100$), nor a significant interaction effect of task × ROI ($F(1.623,17.854) = 1.165, p = 0.324, \omega^2 = 0.013$). The results suggest that the presaccadic potentials differ between the main and memory tasks, albeit to a lesser extent than between the main and visual saccade tasks since potentials for different tasks do not vary across ROIs.

For comparative reasons, an additional analysis between the main and visual saccade task using only 12 participants was conducted, showing similar results as the analysis using all participants. However, the topographic map for the visual task shows slightly more pronounced clusters, this time bilaterally as well as across the midline in parietal electrodes, while still maintaining the left-lateralized centro-frontal cluster (see Figure S7 D). No significant main effect of task was observed ($F(1,11) = 4.233, p = 0.064$, $\omega^2 = 0.084$), but a significant interaction effect of task × ROI was found ($F(1,11) = 8.629, p = 0014, \omega^2 = 0.342$). Post hoc paired t-tests revealed a significant difference for the frontal ROI between the main and visual saccade task ($p = 0.025$), but not for the parietal ROI ($p = 0.211$). There was also no significant difference found for the main task between frontal and parietal ROI ($p = 0.072$). These findings suggest that task-related differences in presaccadic potentials are more pronounced in the frontal regions when comparing the main and visual saccade tasks. The smaller sample size may have contributed to the lack of significant main effect of the task. In contrast, the memory task showed a significant main effect of task, indicating overall task differences, but the lack of a significant interaction effect suggests these differences are consistent across both frontal and parietal regions. This implies that the frontal region is particularly sensitive to task differences in the visual saccade task, while the memory task affects both regions more uniformly. Specifically, the frontal region in the memory task is more similar to the main task, while for the visual task, the frontal region shows a distinct difference compared to the main task. The parietal regions are not significantly different in either task, although the visual task shows some variability indicated by the task × ROI interaction effect. The use of only 12 participants allows for direct comparison between the main, memory, and visual saccade tasks, but may not capture the full variability seen in a larger sample. However, it does clearly highlight the sensitivity and relevance of the frontal regions to task conditions.

A subsequent direct comparison between visual and memory saccade task, including only 12 participants, was conducted (see Figure S7 A-E). The statistical analysis revealed no significant main effect for task ($F(1,11) = 0.026, p = 0.874$, $\omega^2 = 0.000$) or ROI ($F(1,11) = 2.254, p = 0.161$, $\omega^2 = 0.074$). There was also no significant interaction effect of task × ROI ($F(1,11) = 3.341, p = 0.095, \omega^2 = 0.137$). Similarly, repeated measures ANOVA with separate ROIs as factors did not reveal any significant effect. Bayesian analysis revealed a $BF_{10} = 0.686$ for the frontal ROI and $BF_{10} = 1.303$ for the parietal ROI, further indicating largely comparable neural responses between tasks, though the evidence remains inconclusive.

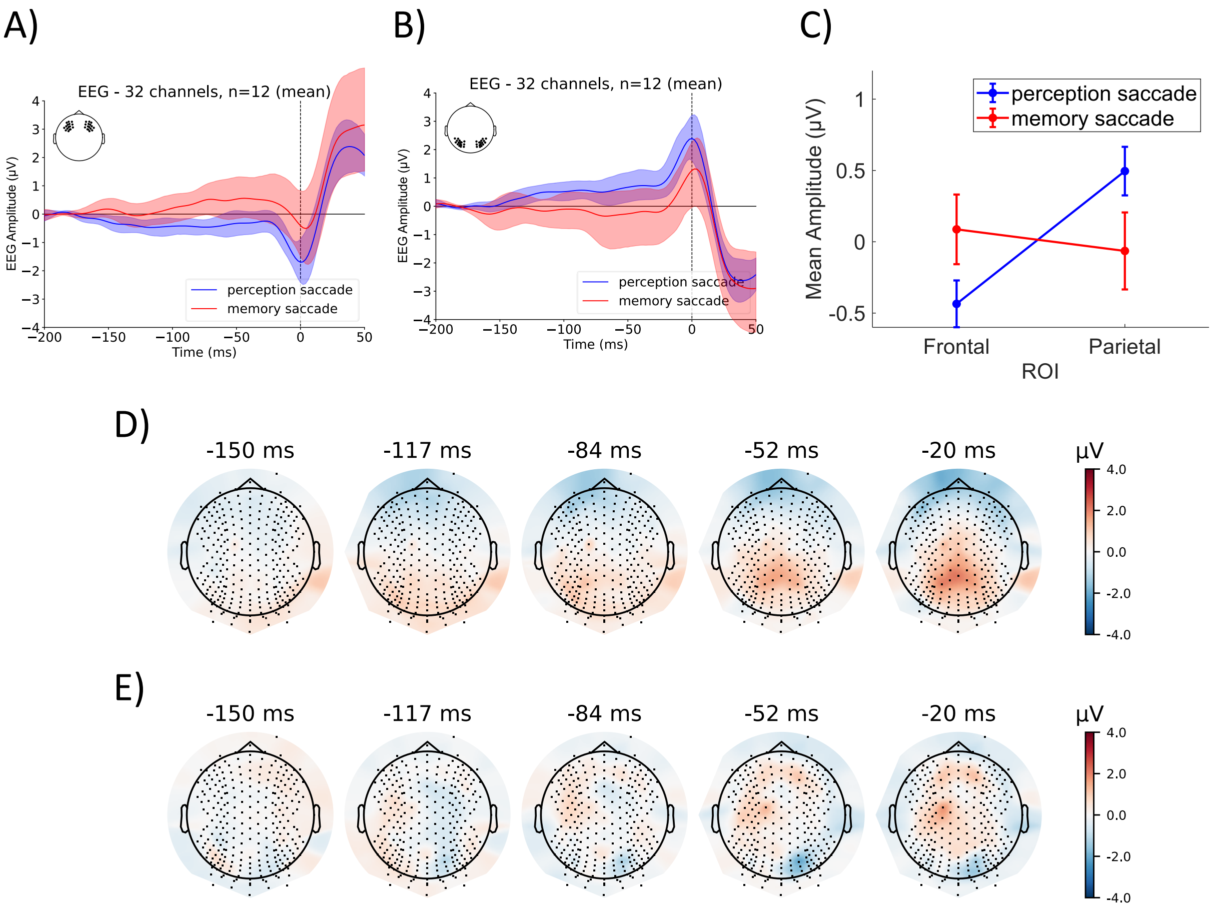


Figure S7. EEG potentials time-locked to visual and memory saccades. 0 ms indicates saccade onset (n = 12). Grand average ERPs with shaded areas representing 95% confidence intervals are shown across A) frontal ROI and B) parietal ROI. C) Mean amplitudes averaged from  150 ms to 20 ms are shown for visual and memory saccade task across frontal and parietal ROIs with error bars indicating the standard error of mean. Topographic maps of presaccadic potentials at selected time points (150 ms to  20 ms) are shown for D) visual and E) memory saccade task.

**Examination of main task conditions**

Moreover, to gain further insights into factors within the main task, additional analyses were conducted. First, the fixed gaze condition was compared to the free gaze condition within the main task (see Figure S8 A-C). The comparison using an ANOVA yielded no significant main $(F(1{,}56) = 0.006, p = 0.939, \omega^2 = 0.000)$ or interaction effect $(F(1{,}56) = 1.116, p =$

$0.295, \omega^2 = 0.001$). The topographic map also revealed no prominent clusters or differences between gaze conditions (see Figure S8 D, E).

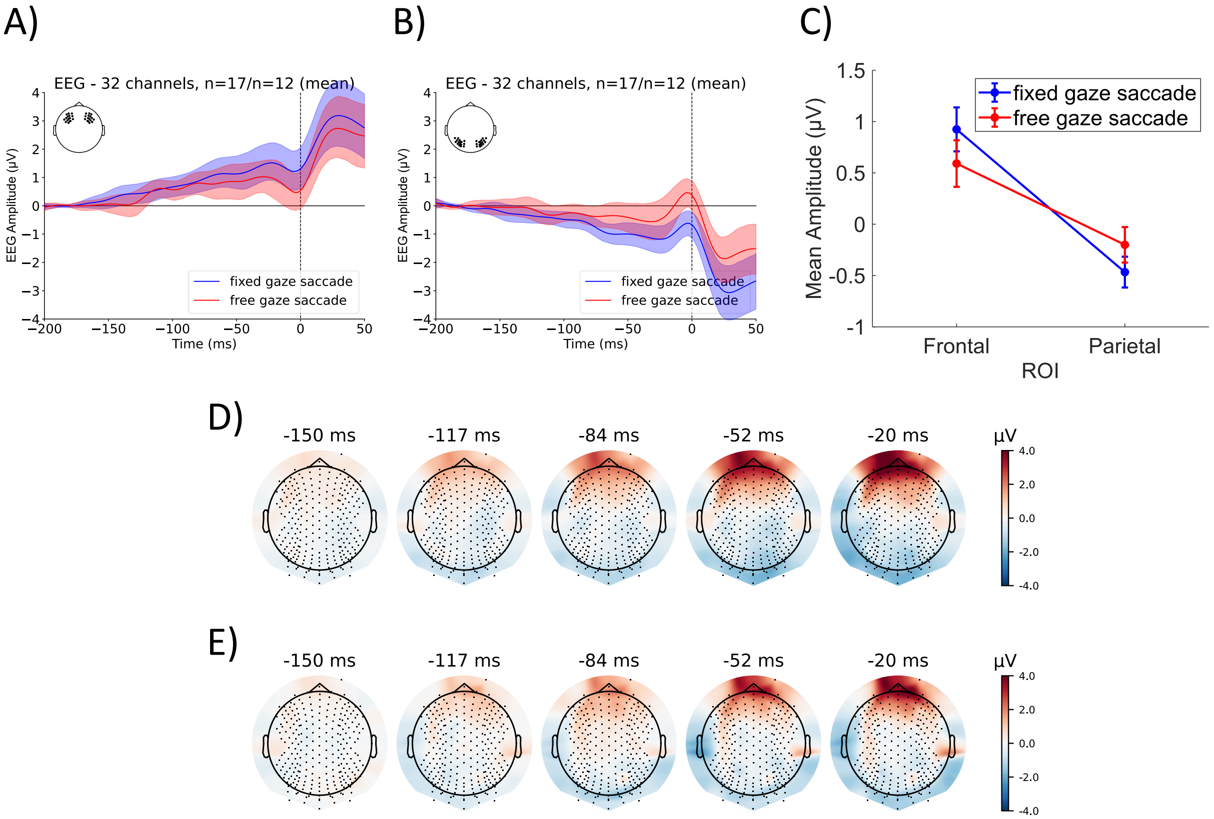


Figure S8. EEG potentials time-locked to saccades during fixed and free gaze conditions. 0 ms indicates saccade onset (n = 17/n=12). Grand average ERPs with shaded areas representing 95% confidence intervals are shown across A) frontal ROI and B) parietal ROI. C) Mean amplitudes averaged from 150 ms to 20 ms are shown for fixed and free gaze conditions across frontal and parietal ROIs with error bars indicating the standard error of mean. Topographic maps of presaccadic potentials at selected time points (150 ms to 20 ms) are shown for D) fixed gaze condition and E) free gaze condition.

Similarly, potentials in the main task for open compared to structured questions were analyzed with results shown in Figure S9 A-C. The open question condition showed a slightly less negative parietal potential than the structured question condition. Nonetheless, repeated measures ANOVA results showed no significant main ($F(1,28) = 1.279, p = 0.268, \omega^2 = 0.003$) or interaction effect $(F(1,28) = 9.668 \cdot 10^{-4}, p = 0.975,\ \omega^2 = 0.000)$ related to the type of question.

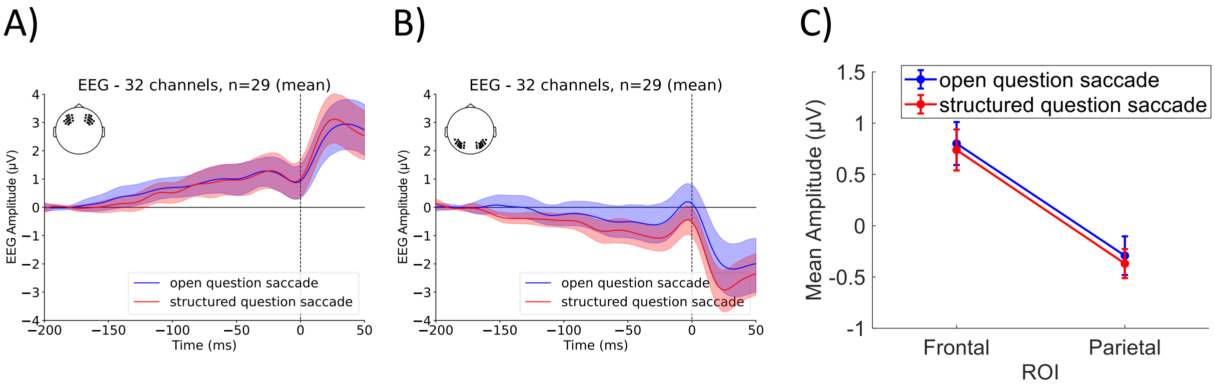


Figure S9. EEG potentials time-locked to saccades during open and structured questions. 0 ms indicates saccade onset (n = 29). Grand average ERPs with shaded areas representing 95% confidence intervals are shown across A) frontal ROI and B) parietal ROI. C) Mean amplitudes averaged from 150 ms to 20 ms are shown for open and structured question types across frontal and parietal ROIs with error bars indicating the standard error of mean.

Lastly, potentials during block 1 were compared to block 4. As seen in Figure S10 A-C, block 4 exhibited a more positive frontal potential and a less negative parietal potential compared to block 1. However, repeated measures ANOVA indicated no significant main $(F(1,28) = 1.753, p = 0.196,\ \omega^2 = 0.013)$ or interaction effect $(F(1,28) = 3.772, p = 0.062, \omega^2 = 0.055)$ related to question block.

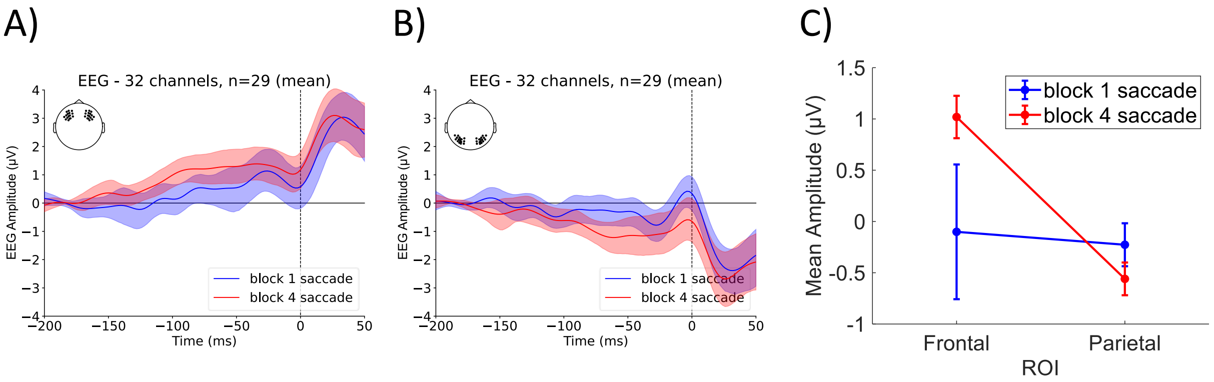


Figure S10. EEG potentials time-locked to saccades during block 1 and block 4 question. 0 ms indicates saccade onset (n = 29). Grand average ERPs with shaded areas representing 95% confidence intervals are shown across A) frontal ROI and B) parietal ROI. C) Mean amplitudes averaged from  150 ms to  20 ms are shown for block 1 and block 4 questions across frontal and parietal ROIs with error bars indicating the standard error of mean.

**Spatial saccade distribution**

For the analysis of saccade directions, since the block factor was non-significant in the analysis, it was excluded to reduce model complexity. To assess saccade direction during cognitive engagement, a GLMM was used on normalized saccade counts with the fixed factors of direction, condition type (fixed/free), and question type (open/structured). Subjects were included as the random effects grouping factor.

Figure S11 A-C shows the overall normalized saccade counts for the direction factor alone for the visual field. An increase in saccade counts in quadrants 1 and 3 can be observed, with slightly higher saccade counts for quadrant 3. When considering the direction factor in combination with condition type and question type, no significant variations were found.

For the quadrant analysis within the visual field, a significant main effect of the direction factor was found $\chi^2 = 12.013, p = 0.007$). Post hoc Wilcoxon signed-rank tests revealed a significant difference between quadrant 1 and 4 ($p = 0.001$), quadrant 2 and 3 ($p < 0.001$), and quadrant 3 and 4 was found ($p < 0.001$). Additionally, significant differences between quadrant 1 and 2 ($p = 0.027$) and quadrant 1 and 3 ($p = 0.031$) were found. No significant difference was found between quadrant 2 and 4 ($p = 0.391$), indicating that there is no difference in saccade counts to the top compared to the bottom. However, there are more saccades to the left than to the right, and overall, more horizontal than vertical saccades. However, ERP analysis did not show lateralized effects, suggesting that while the behavioral data indicate a directional bias, this asymmetry is not translated to the neural activity at the level analyzed.

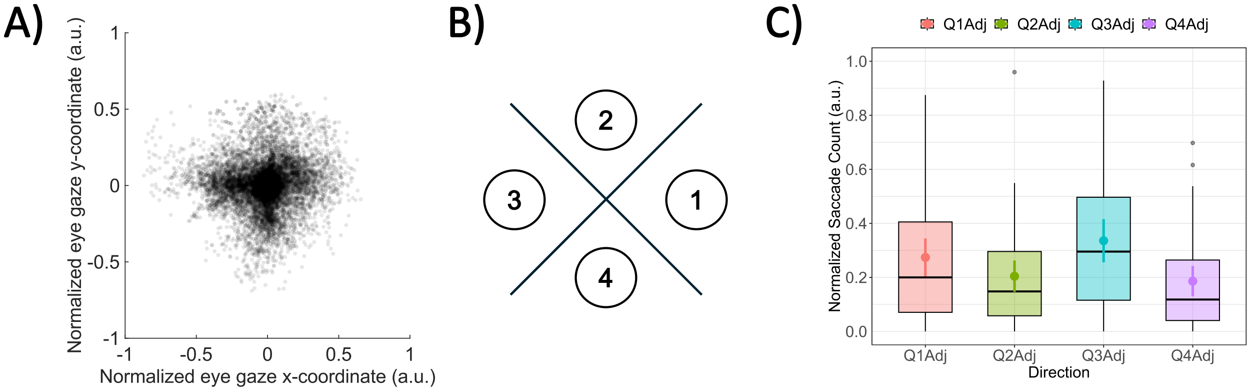


Figure S11. A) Scatter plot of normalized eye gaze coordinates of saccade offsets with participants' centroids subtracted. B) Classification of saccade distributions by saccade-offsets. C) Normalized saccade counts across saccade offset quadrants (Q1Adj–Q4Adj). Box plots show the IQR with medians as horizontal lines. Mean values with 95% confidence intervals are marked as colored points, and outliers as individual points. Only known response trials were included (n = 45).

**Source localization of presaccadic cortical activation**

Main Task

In the main task, source localization revealed broad and intense bilateral activation, with a dominant focus in the right hemisphere, particularly in the superior and middle frontal cortices, extending into the orbitofrontal cortex and precentral gyrus (see Figure S13 A, B). This reflects significant engagement of higher-order cognitive control and motor planning, consistent with the ERP findings that showed diffuse frontal activity. The lack of distinct clusters in motor planning areas like the FEF and DLPFC suggests a reliance on general cognitive control processes rather than precise motor programming. The t-values observed, peaking at $t = 9.3$, were significantly greater than those observed in the control tasks, indicating increased cognitive activity during non-visual saccades.

In the small sample size a cluster at the vertex could be identified which could indicate FEF involvement, however this is not in alignment with ERP results and also not visible in the larger sample size, thus hindering more implications to be made from the finding.

Parietal involvement in the main task was notably absent, which aligns with the ERP results that showed minimal parietal activity and predominant frontal cortex engagement. This reinforces the conclusion that the main task relies on cognitive control mechanisms. However, it is important to note that while we observe significant frontal activation, the lack of identified subcortical structures like the SC limits the ability to fully confirm that the saccades in the main task are triggered by broad frontal activation indirectly projecting to SC. This suggests that further experimentation and refined analysis are needed to pinpoint the precise origin of the activity in the main task, as current methodological approaches may obscure these details.

Visual saccade task

In the visual saccade task, source localization for both the 12-participant and 29-participant groups revealed localized frontal and parietal activation, particularly in the superior parietal lobule and precentral gyrus (see Figure S13 A, B). This corresponds well with the ERP clusters observed in the visual saccade task, possibly reflecting the involvement of the FEF, which plays a key role in saccade programming. There is also evidence of parietal activity spreading from subcortical structures to more superficial cortical regions, aligning with the ERP clusters that suggested a combination of broader PPC involvement, as well as localized PEF involvement in perceptual saccades.

Occipital activity was also present in source localization, reflecting the processing of visual input necessary for perceptual saccades. The parietal and occipital engagement reflects the sensory-spatial integration necessary for visually-guided saccades, which contrasts with the frontal dominance seen in the main task.

Memory saccade task

In the memory saccade task, source localization revealed more localized frontal activation along the midline, but there was also activity spreading to more superficial cortical frontal regions (see Figure S13 A, B). The localized frontal activation likely corresponds to the DLPFC, which was also suggested by the ERP clusters seen in the memory task. Also, the involvement of deeper subcortical involvement can be suggested from source localization, possibly involving the SC, similar to the results seen in the visual saccade task. Moreover, activity in the anterior cingulate cortex could be identified, aligning with previous source localization studies that have linked this region to presaccadic activity (McDowell et al., 2008; Richards, 2013). This pattern of engagement reflects the task's reliance on memory retrieval and voluntary saccade control, as higher cognitive demand triggers deeper and more localized frontal activity.

Interestingly, anterior cingulate cortex (ACC) activation was also identified in the main task for both sample sizes. This finding is notable, as the literature suggests that the FEF (FEFs) and the ACC are commonly coactivated during saccade tasks (Babapoor-Farrokhran et al., 2017; Koval et al., 2014). Given the close functional relationship between these regions, it raises the possibility that the broad activation observed in the main task, particularly involving the ACC, could be contributing to the generation of saccades through indirect pathways. This coactivation supports the idea that even in the absence of specific motor programming, such as in the FEF, cognitive control regions like the ACC could still engage saccadic mechanisms. This, however, remains highly hypothetical and would require much closer analysis to confirm. While the literature does suggest close functional connections between the ACC, FEF, and DLPFC, the exact mechanism by which broad activation in these regions could lead to saccade generation in the main task remains uncertain. Further investigations, with more refined experimental controls, are necessary to clarify whether this coactivation plays a direct role in triggering saccadic movements.

Methodological limitations

When comparing the 29-participant group to the 12-participant group, slight differences in lateralization were observed. For the main task, the larger group exhibited more right-lateralized activation, while the smaller group showed more left-lateralized activity. However, these differences should not be over-interpreted due to methodological limitations, such as downsampling to 64 electrodes and the use of a template head model. These variations suggest that the source localization results should be viewed in terms of broad distributions rather than precise localizations, as the differences between the two groups highlight potential limitations in spatial precision.

The overall patterns across groups, however, remain consistent in that the main task shows broad, diffuse frontal activation, while the control tasks show more focused and localized activation in both frontal and parietal regions.

Additionally, it is important to note that the source localization model used in this study only provides labels for cortical structures, meaning that any involvement of subcortical regions, such as the SC, must be inferred based on broader cortical activation patterns rather than directly identified.

Distinct neural mechanisms for visual vs. non-visual saccades

The source localization results enabled us to confirm the previous inference that the main task is substantially different from both the visual and memory saccade tasks, as the latter two show much more localized activity. This is particularly evident in the clear frontal and parietal activation patterns seen in the control tasks, which align closely with ERP findings and the existing literature. The ERP data from the control tasks align well with established research, demonstrating the expected engagement of sensory-spatial integration and motor planning mechanisms. In contrast, the main task presents a different pattern, with broad and diffuse frontal activation, confirming our earlier observations from ERP results that the main task does not follow typical saccadic programming patterns.

The key finding is that while source localization confirms the distinctions between the main task and control tasks, the main task presents a unique challenge. We are unable to directly pinpoint specific regions responsible for saccade generation in the main task, as both source localization and ERP results show broad frontal engagement without clear focal clusters. The lack of precise localization, coupled with the absence of identifiable subcortical structures such as the SC, limits our ability to fully support the hypothesis that frontal regions indirectly contribute to saccade generation during non-visual tasks. Although the identification of the anterior cingulate cortex (ACC) aligns with literature on its coactivation with the FEF during cognitive saccade tasks, this connection remains speculative. Further analysis is needed to explore the functional interactions between these regions in non-visual saccades.

These results underscore the need for more controlled main task conditions and future experiments with fewer confounding factors. Subtle effects may be masked by the exploratory nature of the current setup, preventing us from conclusively identifying the neural origins of non-visual saccades. More refined analyses and controlled experimental designs are required to fully understand the mechanisms behind these diffuse activations, particularly in non-visual saccades, and to confirm how these processes differ from visual and memory-guided saccades.

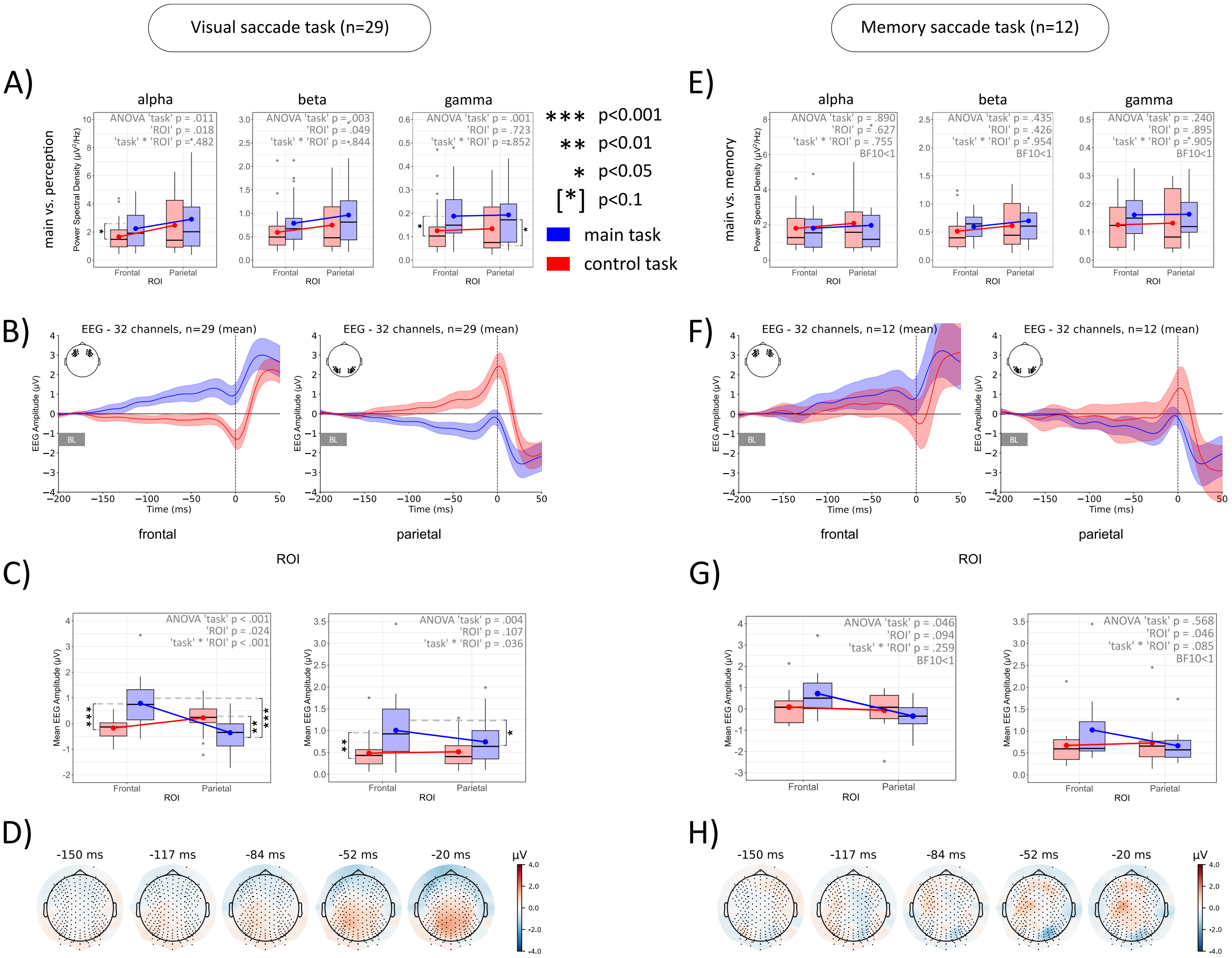


Figure S12. Comparison of the main task vs. visual and memory saccade control tasks. A) Modulation of alpha, beta and gamma frequency across tasks. B) ERPs for VST and the MST tasks; C) Average ERP amplitudes across tasks. D) Topographic maps of presaccadic activity across both control tasks. Error bars denote upper and lower quartiles, boxes denote interquartile ranges, solid horizontal bars denote medians.

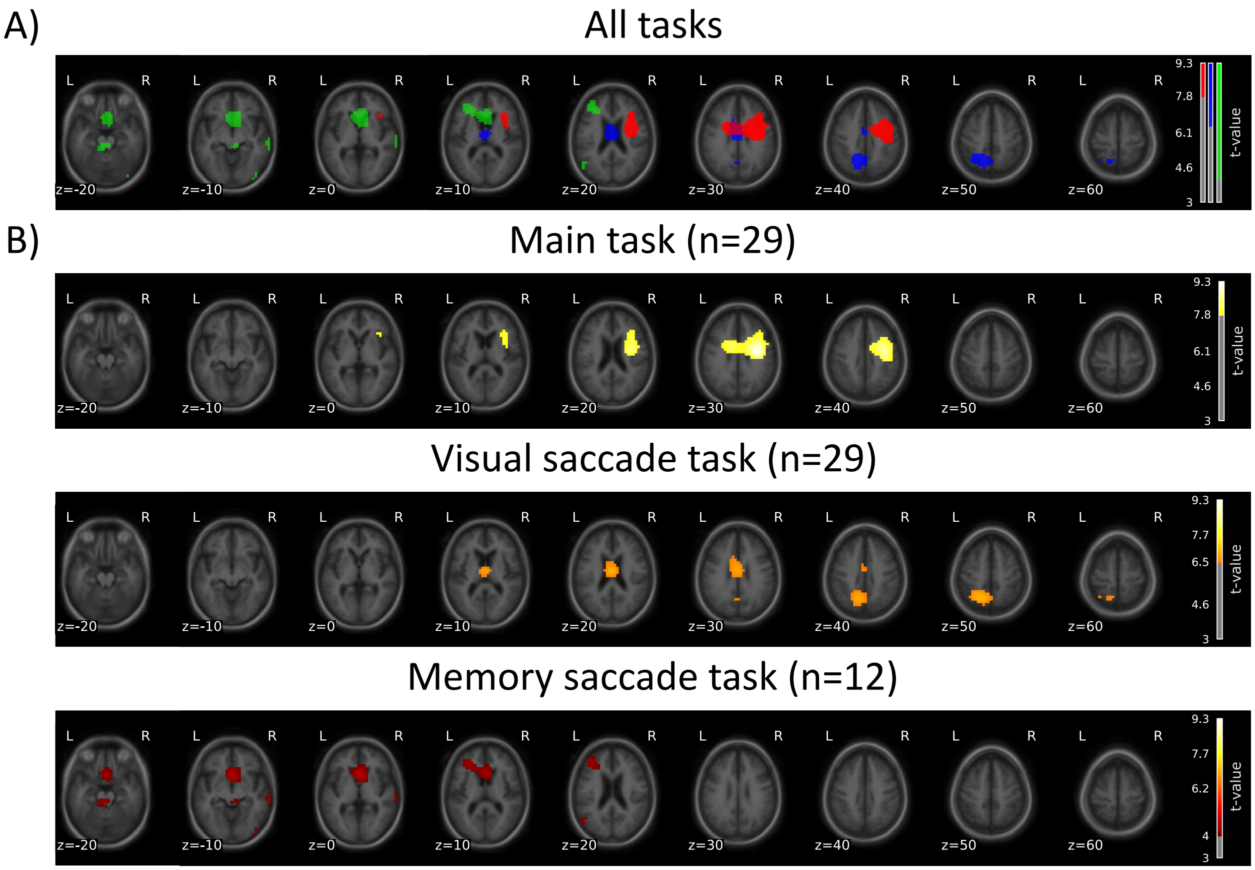


Figure S13. Axial slices of current sources for the main task (red), VST (blue), and MST (green), shown separately (A) and combined (B).

Table S5. Overview of participant exclusions across analysis steps based on applied criteria.

| Analysis Step | Reason for Exclusion | N Excluded | N Remaining |
|---|---|---|---|
| **1. Behavioral** | <30 trials post-processing, incomplete dataset | 8 | 45 |
| | main sequence, SD/RMS outliers, visual inspection | 4 | |
| **2. EEG Main Analysis** | invalid control task eye data: incomplete dataset, visual inspection | 3 | 29 |
| | incomplete EEG dataset, trigger loss | 8 | |
| | <40 main task trials post-processing | 4 | |
| | <40 control task trials post-processing | 6 | |
